# Widely dispersed clonal expansion of multi-fungicide-resistant *Aspergillus fumigatus* limits genomic epidemiology prospects

**DOI:** 10.1101/2024.07.29.605539

**Authors:** Eveline Snelders, Brandi N. Celia-Sanchez, Ymke C. Nederlof, Jianhua Zhang, Hylke H. Kortenbosch, Bas J. Zwaan, Marlou Tehupeiory-Kooreman, Alejandra Giraldo-López, Karin van Dijk, Li Wang, Marin T. Brewer, Michelle Momany, Ben Auxier, Paul E. Verweij

## Abstract

**Background:** *Aspergillus fumigatus* is a ubiquitous fungus that causes a range of diseases in animals, including humans. The most lethal manifestation is invasive aspergillosis for which treatment relies on triazoles. Triazole-resistant *A. fumigatus* can be recovered from decaying plant material and so-called hotspots containing triazole fungicide residues. Although observations have shown clonal isolates between the environment and clinical samples, a direct link between a specific environment and cases of triazole-resistant invasive aspergillus disease in an individual patient has not yet been demonstrated.

**Methods:** To understand where patients acquire *A. fumigatus* isolates causing disease, we used a genomic epidemiology approach with 157 Dutch *A. fumigatus* isolates, based on whole genome sequencing. Isolates were from three well-characterized environmental hotspots and two hospitals between 2016 and 2019.

**Findings:** In the Dutch dataset, *A. fumigatus* isolates from six patients showed near-identical genomes compared to five environmental isolates. One environmental isolate matched three probable cases of triazole-resistant invasive aspergillosis, including one fatal case. Patient isolates were recovered up to 34 months later than near-identical environmental isolates. Comparison to over 1·2K global publicly available *A. fumigatus* genomes showed hundreds of clonal groups spread across three continents. In addition, finding variants associated with resistance to non-triazole fungicides such as benzimidazole, succinate dehydrogenase inhibitor and quinone outside inhibitor classes, strongly suggests an exposure history to multiple agricultural fungicides in these environmental hotspots.

**Interpretation:** Environmental hotspots represent highly selective habitats for multi-fungicide-resistant *A. fumigatus*, which we can now directly link to probable cases of aspergillus disease, including a triazole-resistant case. However, geographically widely dispersed clonal expansion limits the utility of genomic epidemiology to identify the source of a particular patient’s isolate. Furthermore, reducing a single class of fungicides in agriculture may not effectively reduce resistance selection when other classes are still in use.

**Funding:** NWO.Groen2019.002

**Research in context:** *Evidence before this study:* Triazole fungicides that exhibit activity against *Aspergillus fumigatus* have been shown to be a major source of resistant aspergillus disease in humans. However, the route of transmission from environmental hotspot to human remains poorly understood. Isolates of *A. fumigatus* can be recovered from both environmental samples and clinical specimens that harbour the same resistance signature haplotypes, e.g., TR_34_/L98H and TR_46_/Y121F/T289A, in the *cyp*51A-gene. We searched the literature for evidence using high resolution whole genome sequencing (WGS) to link environmental isolates to human infection. We searched PubMed for articles using the search terms ‘*Aspergillus fumigatus*’ AND ‘azole resistance’ AND ‘whole genome sequencing’ on 15 April 2024. This search retrieved 32 articles describing different evolutionary routes to select for triazole-resistant *A. fumigatus* or population structure of whole-genome sequenced isolates. Twenty-six articles used whole-genome sequencing, but none focused on identifying clonal groups to identify direct cases of transmission between the environment and clinical cases of aspergillus disease. By using the additional search term ‘transmission’, no other records were retrieved.

*Added value of this study:* Our study links triazole-resistant *A. fumigatus* isolates cultured from three environmental hotspots to cases of aspergillus disease in two hospitals in the Netherlands. Genome comparisons of isolates from environmental hotspots and patients showed multiple near-identical linked genotypes, consistent with a route of transmission from the environment to patients. Although a naïve expectation may be a higher probability of matches of the hotspots located in the northwest of the Netherlands with the hospital located in the west of the country, in fact, more patient isolates from the far southeast were linked to the hotspots. Integrating the Dutch data set into a global data set showed 205 clonal groups spread across the Netherlands, Germany, the United Kingdom (UK), the United States of America (USA) and Japan. Our demonstration of a large number of geographically dispersed clonal groups suggests that current sampling is insufficient to definitively identify the source of an individual patient’s infection. A genetically highly diverse population combined with a wide global distribution of clones can make it impossible to definitively identify the source of an individual patient’s infection even with much more sampling.

*Implications of all the available evidence:* Our study provides evidence that triazole-resistant *A. fumigatus* isolates with multi-fungicide resistance profiles cause aspergillus disease in at-risk patients and may contribute to treatment failure and mortality. The risk of infection due to these triazole-resistant isolates is not confined to the geographic vicinity of the environmental hotspot since clonal spread can be detected across great distances. The finding of linked cases without clear transmission routes limits epidemiological studies and underscores the need to better understand the ecology and environmental niches of this fungus. As it is highly unlikely that each patient visited the rural agricultural areas where a hotspot was located, research should address the complex and long-distance transmission routes of resistant isolates, which involves airborne dispersal of conidia or habitats of this fungus outside the agricultural environment. Furthermore, because of the multi-fungicide resistance phenotype of the triazole-resistant *A. fumigatus*, involving several classes of fungicides, reducing one class of fungicides in the environment may not effectively reduce resistance selection. Effective interventions should instead aim to reduce the burden of environmental resistance by modifying environments that currently favour the massive outgrowth of fungicide-resistant *A. fumigatus* to limit the escape of aerial spores from these environmental hotspots.

## Introduction

*Aspergillus fumigatus* is a ubiquitous fungus that causes a range of diseases in animals including humans. The fungus thrives on decaying plant material and produces large numbers of conidia, which following airborne dispersal may be inhaled by humans and subsequently cause pulmonary disease in susceptible hosts. The most lethal manifestation of aspergillus disease is invasive aspergillosis (IA), where invasive growth of the fungus leads to death if left untreated. Susceptible patients include those with a haematological malignancy, solid organ transplant recipients, and hematopoietic stem cell recipients, and critically ill patients with severe influenza and COVID-19 infection.^1–3^ Treatment of IA and other forms of aspergillus disease relies largely on triazoles, including itraconazole (ITR), voriconazole (VOR), posaconazole (POS), and isavuconazole (ISA). The introduction of the triazole class has significantly improved the survival of patients with IA, with a 15% improved survival in patients treated with VOR compared to those receiving treatments based on amphotericin B.^4^ However, the efficacy of triazole therapy is threatened by the emergence of resistance in *A. fumigatus*, observed since the 2000s^5^. Triazole resistance evolves de novo in a minority of cases in-host during therapy, while in parallel exposure to triazole fungicides in the environment has proven to be the main route of resistance selection.^6^ Although not a target pathogen, many agricultural triazole fungicides exhibit activity against *A. fumigatus*.^6^ In the Netherlands high concentrations of triazole-resistant *A. fumigatus* are recovered from decaying plant material that contain residues of triazole fungicides.^7^ These hotspots for resistance selection include flower bulb waste, green waste, wood chippings, and organic waste.^7^ Triazole resistance variants commonly involve changes in the *cyp*51A gene, most notably single nucleotide polymorphisms (SNP) combined with tandem repeats (TR) of different lengths in the gene promoter, e.g. TR_34_ (34-bp tandem repeat duplication) with an L98H nonsynonymous SNP and TR_46_ (46-bp tandem repeat duplication) with Y121F and T289A non-synonymous SNPs, also denoted as TR_34_ or TR_46_ *cyp*51A haplotypes.^5,8^ These variants confer resistance to agricultural triazoles as well as cross-resistance to medical triazoles.^9^ Such resistance presents several challenges for the management of patients with IA, including drug-resistant disease in triazole-naïve patients and patients with mixed triazole-susceptible and triazole-resistant infection.^5,8,10^ A retrospective cohort study indicated that the 90-day mortality rate was 25% higher in patients with VOR-resistant IA compared with patients with VOR-susceptible infections, when treated with VOR mono-therapy.^11^

The growing availability of whole genome sequencing (WGS) of *A. fumigatus* has revealed various important new insights, such as new variants associated with antifungal drug resistance.^12^ Near-identical triazole-resistant haplotypes of environmental origin and clinical samples of *A. fumigatus* supports a transmission route of isolates from the environment to patients,^13^ confirmed by previous studies based on microsatellite typing.^14,15^ Although these observations support a strong link between selection of environmental resistance and clinical infection, the geographic spread of *A. fumigatus* strains resistant to triazoles remains poorly understood. To date, no direct link of transmission has been established between environmental hotspots and cases of triazole-resistant aspergillus disease.

In this study, we used WGS to investigate the genomic epidemiology of 157 *A. fumigatus* isolates, obtained from three well-characterized close-by environmental hotspots located within a 4 kilometres/2·5 miles radius and two hospitals, one closer to the environmental hotspots (65 kilometres/40 miles) and the other at a larger distance (150 kilometres/93 miles). We used genomic data to investigate the population structure and genomic variation associated with phenotypic resistance to medical triazoles. We also tested antifungal susceptibilities of a subset of isolates to medical and agricultural antifungal compounds including triazoles as well as non-triazole fungicides with other modes of action. We place this in a global context by comparing the Dutch *A. fumigatus* genomes from this study with those publicly available and collected world-wide.

## Methods

### Study design

Environmental and clinical isolates of *A. fumigatus*, identified based on growth at 48°C and macroscopic colony morphology, were selected to attempt a 1:1 match of the year of isolation, the *cyp*51A genotype, and the triazole susceptibility phenotype (Supplemental Material Figure S1). The environmental isolates were recovered from samples of plant waste material collected in a previous study^16^ and included three flower bulb farms (A, B, and C) in the northwest of the Netherlands, hereafter indicated as “environmental hotspots” for triazole resistance. Furthermore, we included six triazole-resistant *A. fumigatus* isolates cultured from water samples in 2015 from municipal water purification plants in the Netherlands. The clinical isolates of *A. fumigatus* were selected from two Dutch University Medical Centres: the Radboud University Medical Centre (RUMC) in Nijmegen and the Amsterdam University Medical Centre (AUMC). For the years 2016-2019, clinical microbiological records of the two hospitals involved in this study were screened for patients with a positive culture result for *A. fumigatus*. The medical records were then investigated to determine whether the patient had aspergillus disease, including IA and chronic pulmonary aspergillosis (CPA). For aspergillus disease classification, international consensus case definitions were used, including the European Organisation for Research and Treatment of Cancer (EORTC)/Mycosis Study Group Education and Research Consortium (MSGERC) definitions, the AspICU definitions, the influenza-associated pulmonary aspergillosis (IAPA) expert case definition, and the 2020 ECMM/ISHAM COVID-19 associated pulmonary aspergillosis (CAPA) case definition. For CPA patients, the ECMM/ISHAM/ERC case definitions were used.

### Procedures

Selected clinical *A. fumigatus* cultures were tested for triazole susceptibility using an agar-based method (VIPcheck^TM^, MediaProducts, The Netherlands), which employs RPMI-1640 agar supplemented with ITR, VOR, POS. For *A. fumigatus* isolates with a triazole-resistant phenotype, the complete *cyp*51A gene including its promoter was analysed by PCR amplification and Sanger sequencing. All clinical isolates were stored at -70°C in 10% glycerol. The 85 environmental *A. fumigatus* isolates were cultured from the above mentioned flower bulb farms sampled on 25 unique days between July 1, 2016 and July 21, 2017.^16^ The additional six isolates from water samples were all cultured in 2015. Suspensions were incubated on Flamingo agar at 48°C for five days at high density.^17^ Subsequently, by using a 100-needle replicator, colonies were transferred to Malt Extract Agar (MEA) plates with and without triazoles (ITR 0·5 mg/L and 4 mg/L, VOR 0·5 mg/L, and POS 2 mg/L). Colonies of varying triazole susceptibilities were subcultured from the MEA plate without triazoles. All environmental *A. fumigatus* isolates were tested for triazole resistance (ITR, VOR and POS) using VIPcheck^TM^ testing^18^ and the *cyp*51A promoter region and coding gene were analysed for SNPs and indels by Sanger sequencing and compared to AFUA_4G06890.^16^ Subsequently, isolates were single spore passaged and frozen at -70°C in 20% glycerol. Finally, isolates were selected for WGS according to triazole resistance phenotype and *cyp*51A haplotype, and DNA was extracted as previously described.^19^ DNA was extracted from a total of 170 clinical and environmental isolates for sequencing at a commercial provider (Baseclear, the Netherlands), DNA of all clinical isolates was extracted at the RUMC and DNA of all environmental isolates was extracted at Wageningen University & Research. Genomic libraries were prepared using Nextera Flex and sequenced on an Illumina NovaSeq 6000, 250-bp flowcell, with a minimum of 3 Gb of data per sample by a commercial provider (Baseclear, The Netherlands). Four genomes of separately sequenced clinical *A. fumigatus* isolates were added to the analysis. In addition to the Dutch WGS, an extended analysis was performed with 1231 global *A. fumigatus* isolates (Germany, United Kingdom, Japan, and the USA) publicly available through NCBI.^12,13,20–26^

### Data analysis

A detailed description of bioinformatics methods including the code supporting the analysis of the mapping, variant calling, phylogenetic network analysis, and population differentiation analysis can be found in Supplemental Methods and https://github.com/fungalsnelderslab/2024_Dutch_fumigatus. Sequences were deposited in the European Nucleotide Archive (ENA) under project PRJEB73793.

## Results

### Near identical clinical and environmental *A. fumigatus* isolates

To investigate whether *A. fumigatus* isolates from the Netherlands are clonally related in this study, we calculated the distribution of pairwise genetic distances between isolates. Based on histograms (Supplemental Materials Figure S2), a cut-off distance of 0.998 was used, resulting in 22 clonal groups among the Dutch isolates. These clonal groups showed a mean of 76.1 (min: 17; max: 146) high-confidence variants across the whole genome. Of these clonal groups, five were composed of both clinical and environmental isolates, with nine groups made up of only environmental isolates and eight with only clinical isolates. In total, isolates from six Dutch patients matched to five Dutch environmental hotspot isolates (Table 1). Isolates from three patients with probable IA due to triazole-resistant *A. fumigatus* harbouring the TR_34_/L98H *cyp*51A haplotype were matched to isolates from agricultural plant waste material in the northwest of the Netherlands. The cases occurred over a period of 17 months in RUMC, although one patient (Table 1, clonal group A) had been transferred from a hospital located in the south of the Netherlands (south area of the Limburg province). This patient had been transferred due to respiratory deterioration, and the isolate that underwent WGS was recovered from a bronchoalveolar lavage (BAL) sample that was obtained one day after the transfer. Of the three patients with triazole-resistant IA, one patient with IAPA died, which was considered attributable to IA. A triazole-resistant *A. fumigatus* isolate from a patient with allergic bronchopulmonary aspergillosis (ABPA) and CPA, with a wild-type *cyp*51A sequence, matched wild-type *A. fumigatus* isolates from two environmental hotspots (Table 1; case group B). Two patients from AUMC, that were colonized with *A. fumigatus* TR_46_/Y121F/T289A *cyp*51A haplotype, matched with one isolate cultured from potable water at a site in the northeast of the Netherlands (southwest part of the Drenthe province) (Table 1; case group C) and the other patient isolate matched with one recovered from an environmental hotspot in the northwest of the Netherlands (north area of the Noord-Holland province; Table 1; case group D). All four *A. fumigatus* isolates of patients with aspergillus disease showed multiple triazole phenotypes in subsequent cultures (Table S1 of Supplemental Materials).

**Table 1.**
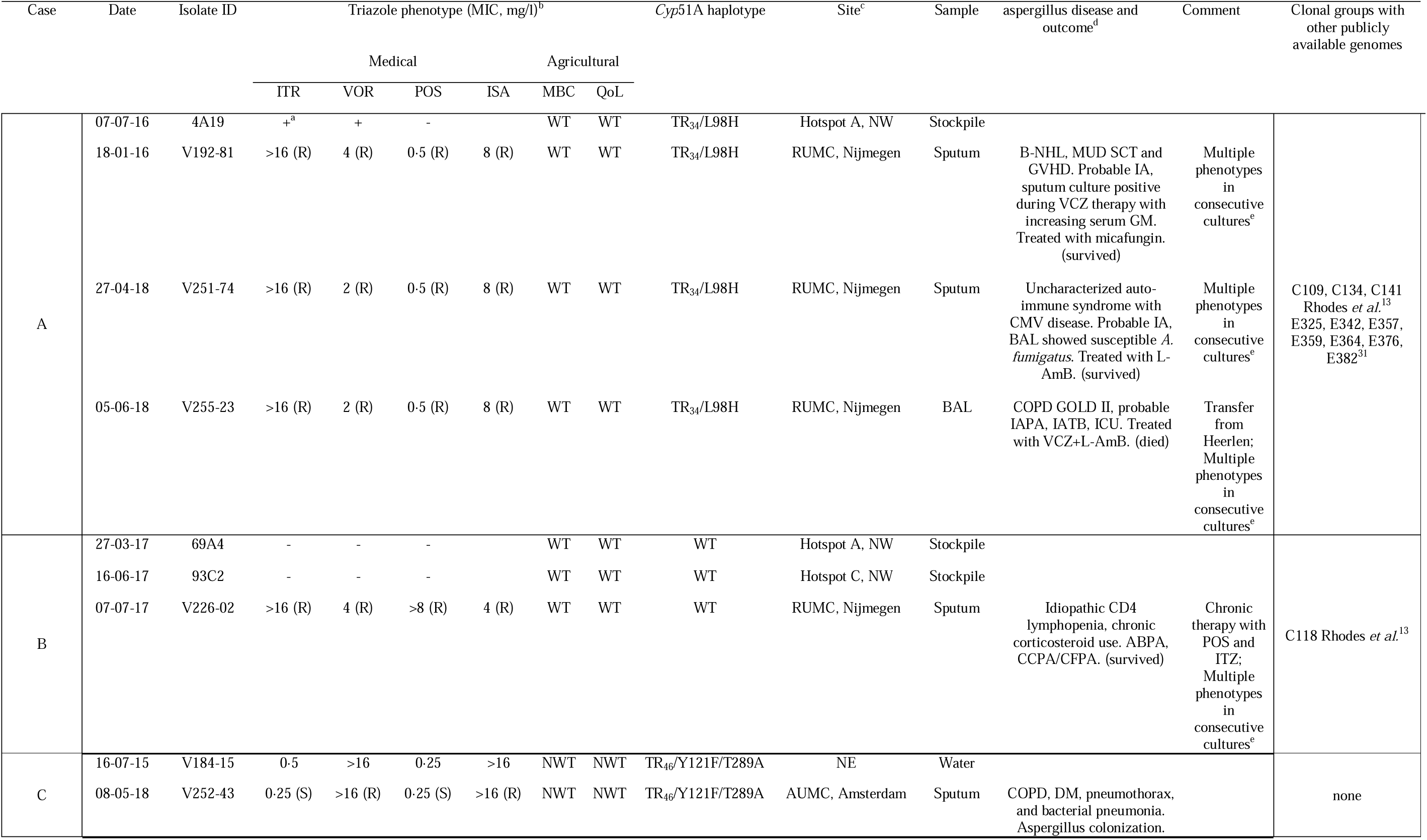
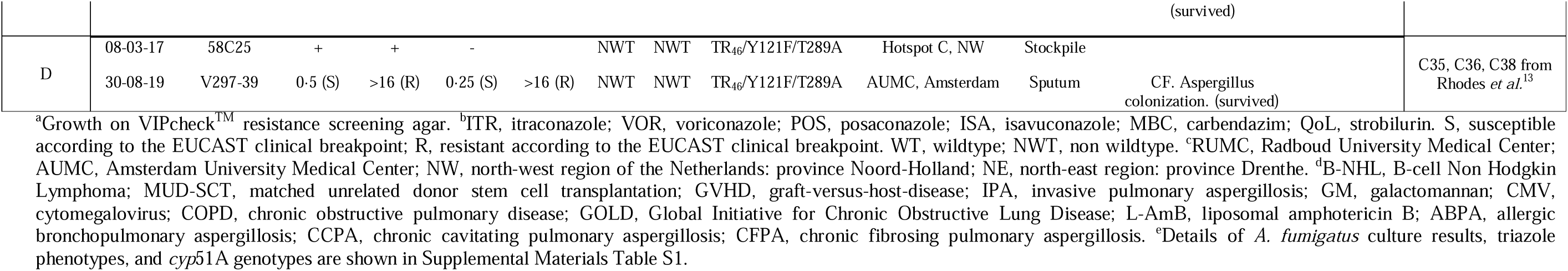
Details of six clinical *A. fumigatus* culture positive cases linked to environmental *A. fumigatus* isolates.

Expanding the same pairwise genetic distance distribution as used on the Dutch samples to analyse the larger global genome dataset, including ∼1·2K publicly available *A. fumigatus* genomes, a total of 205 clonal groups were detected. In Figure 1, all isolates that could clearly be identified as clinical or environmental and with a known location (country) are depicted resulting in a total of 197 clonal groups. Clonal groups limited with clinical and environmental genomes unique to a single country could be identified in this dataset for the USA (n=14), Germany (n=7), UK (n=4) and Netherlands (n=2). A total number of 28 clonal groups were identified across countries between clinical and environmental genomes. Within and across countries between environmental genomes only, 16 clonal groups were detected and five between clinical genomes only.

**Figure 1.**
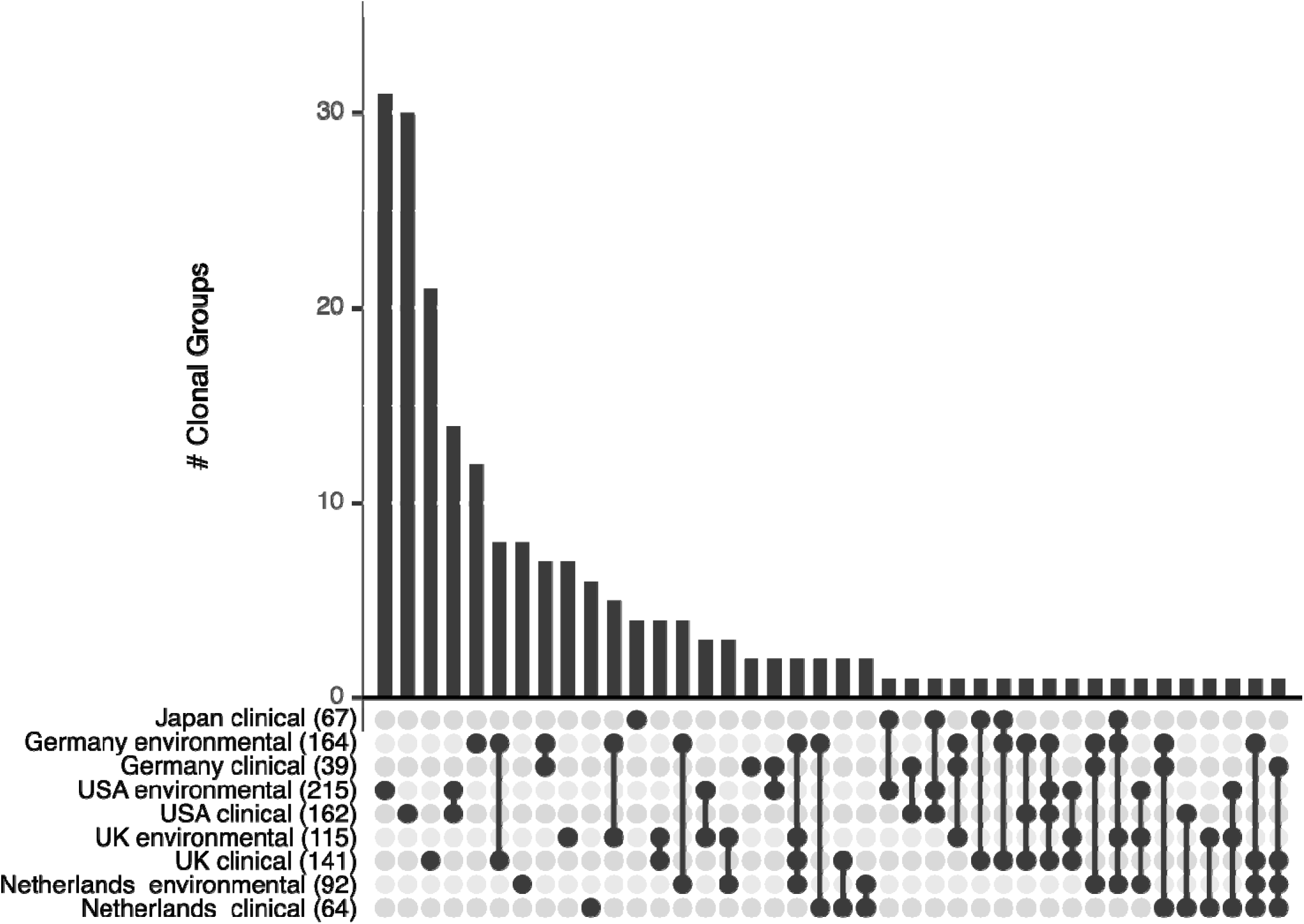
Clonal groups of environmental and clinical *A. fumigatus* isolates. Upset plot indicating 197 unique clonal groups of ∼1·2K global *A. fumigatus* with clearly identified environmental or clinical sources and geographic localisation of Japan,^21^ Netherlands (this study), United Kingdom,^13^ Germany,^26,27^ USA^20,22–25^. Numbers in brackets indicate total isolates from each category. Multiple clonal groups are detected between environmental and clinical isolates within countries or are shared among up to four countries.

### Population genomic analysis

We used principal component analyses (PCA) on the Dutch *A. fumigatus* genomes to visualise variance and, therefore, patterns in the population. No clustering was observed on year of sampling, geographical location, or type of isolate (clinical or environmental) (Supplemental Materials Figure S3). Clustering is, however, seen on TR *cyp*51A haplotypes, where triazole-resistant isolates harbouring the TR_34_, TR_46_, or TR_92_ *cyp*51A haplotypes cluster together in a subset of the space occupied by the isolates with a wild-type *cyp*51A haplotype. To investigate the population structure, an analysis was performed to determine the most likely number of genetic clusters amongst the sampled population, or K value. An exponential decay of the cross-validation value across values of K was observed which strongly indicates the absence of discrete clusters in the *A. fumigatus* population (Supplemental Materials Figure S4). This confirms the finding of the study by Barber *et al.* where multiple (up to seven) clusters were identified in 300 genomes.^12^ Other recent studies^13,21,22,24,25^ however do show discrete clusters, sometimes incorrectly defined as clades. A cluster is a group of isolates grouped based on genetic similarity, irrespective of their evolutionary relationship, so it may or may not reflect common ancestry. A clade, however, is a group of isolates defined by a common ancestor that forms a monophyletic group. A clade is therefore defined entirely on proving common ancestry. As recent studies indicate significant gene flow between *A. fumigatus* populations,^12,13,20,25^ we used a phylogenetic network to analyse relatedness between isolates. Compared to traditional phylogenetic tree analyses, like neighbor joining methods, phylogenetic networks capture evolutionary processes such as recombination, hybridization, and lateral gene transfer. The phylogenetic network of the Dutch isolates (Figure 2a) shows a general separation of the triazole-resistant and triazole-sensitive isolates, although triazole-sensitive isolates are also observed within the triazole-resistant cluster. The multiple paths at the centre of the network indicate ongoing abundant recombination and therefore active sexual reproduction of this fungus, previously detected in multiple studies.^13,24,25,27^ Integrating the Dutch isolates into a larger global data set of ∼1·2K genomes showed a similar pattern (Figure 2b), with a web-like structure of genomic relationships consistent with significant ongoing recombination in the *A. fumigatus* population.^28^

**Figure 2.**
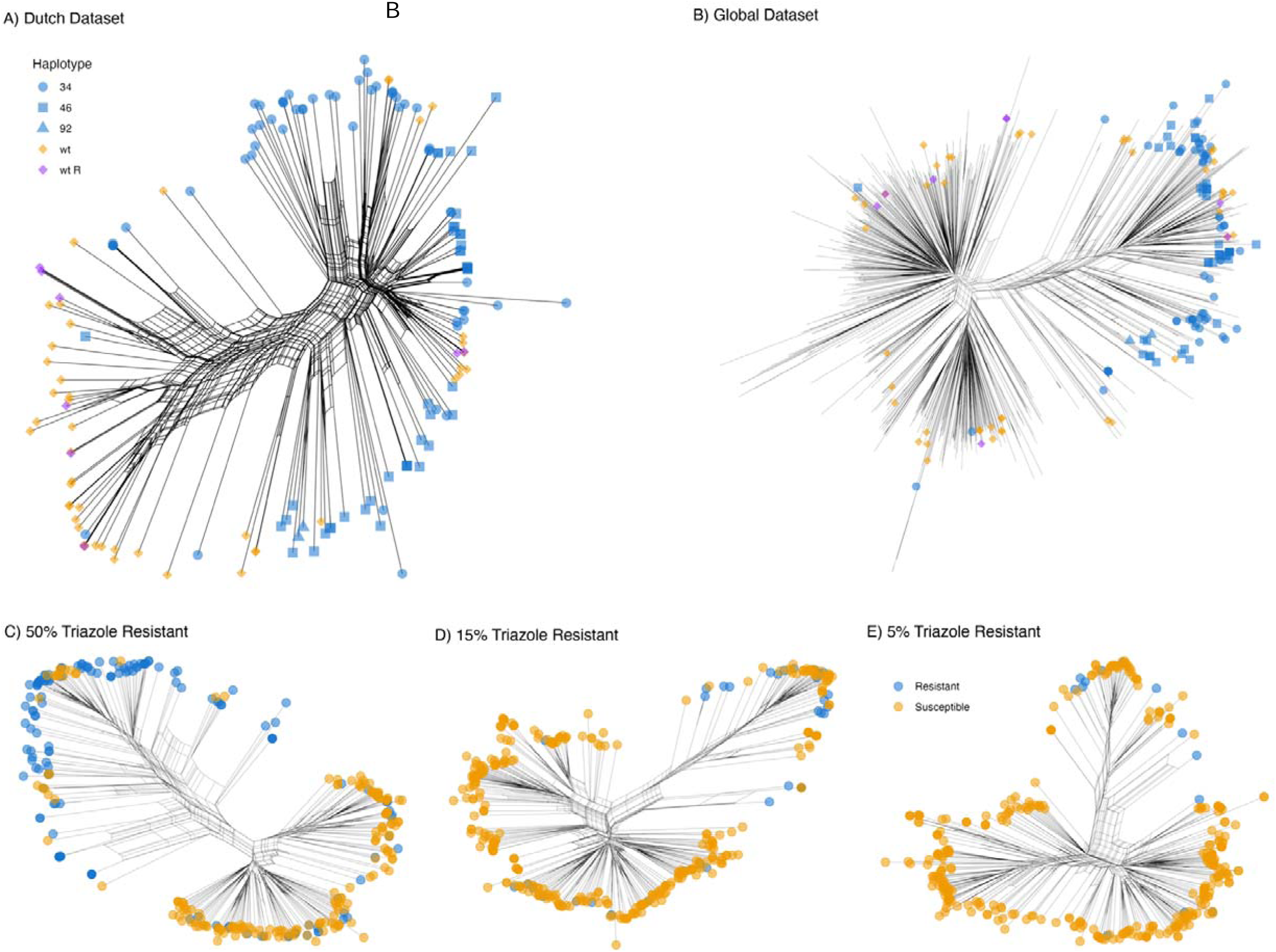
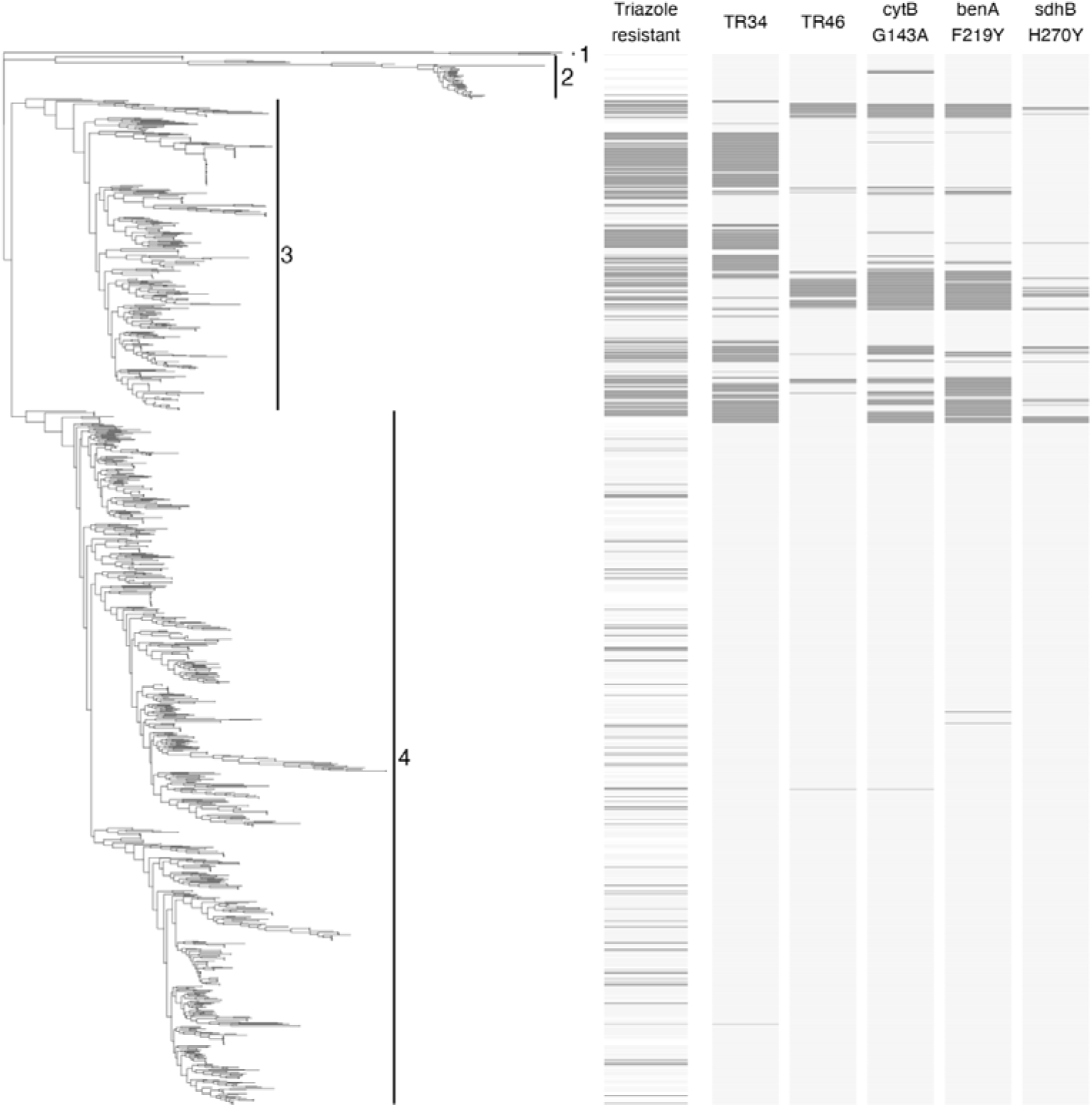
*A. fumigatus* population genomic analysis. a, Phylogenetic network of the 157 Dutch *A. fumigatus* isolates with tips coloured blue; triazole-resistant isolates with a TR *cyp*51A haplotype, in purple; triazole-resistant isolates with a wild type *cyp*51A haplotype, and in orange; triazole-susceptible isolates with a wild type *cyp*51A haplotype. Multiple network paths connecting them indicates ongoing abundant recombination. b, Phylogenetic network of the Dutch (n=157) isolates plus all global (n=1231) genomes, showed a similar pattern with again significant recombination between triazole-resistant and sensitive groups. Note for that tips of global isolates are not coloured to avoid overplotting. c-e, Subsampling of the global *A. fumigatus* genomes in different susceptible to resistant ratios to test the effect of sampling bias on the resulting network structure; tips colored in blue triazole-resistant isolates, and in orange; triazole-susceptible isolates, percentage of triazole-resistant isolates in c; 50% triazole resistance, d; 15% triazole resistance, e; 5% triazole resistance. f, Neighbor joining tree of 157 Dutch (n=157) isolates plus all global (n=1231) genomes. Cluster 1 shows the root of the tree by one *Aspergillus oerlinghausenensis* and four *Aspergillus fischeri* genomes. Cluster 2 shows previously identified distant *A. fumigatus* group of isolates.^25,27^ Cluster 3 shows the oversampled TR *cyp*51A haplotypes together with triazole susceptible wild type *cyp*51A haplotypes. And finally, cluster 4 containing triazole susceptible and triazole resistant mostly non-TR *cyp*51A haplotypes. The panels next to the tree show from left to right; phenotypic triazole susceptibility (based on available metadata), presence of the TR_34_ *cyp*51A, TR_46_ *cyp*51A, *cyt*B^G143A^, *ben*A^F219Y^ and *sdh*B^H270Y^ alleles.

It is important to realise that in many studies, like our own, isolate selection is not random but intentionally oversamples triazole-resistant TR_34_ or TR_46_ *cyp*51A haplotypes, often equal or more than 50% and similar to our study.^25,29^ Since patients and plant waste material both become infected or inoculated by spores from the air, sampling the air should rather act as a representative sample of the population at large. Recent studies showed that triazole resistance of airborne spores of *A. fumigatus* is common but varies between 0 and 10% for itraconazole and voriconazole.^30,31^ Therefore, we subsampled the global *A. fumigatus* genomes in ratios that reflect these airborne resistance levels, to observe the effect on the resulting apparent population structure of triazole-resistant isolates (Figure 2c-e). Using a 50:50 ratio, the current standard analysis, divides the population in the phylogenetic network tree. However, in the 15% and 5% triazole resistance subsampling’s this division is lost, and triazole-resistant genomes are grouped together with triazole-susceptible genomes, not separated from them.

To compare our study with previously generated neighbor joining trees, we also constructed one with the Dutch genomes of this study and the ∼1·2K global genomes (Figure 2f). In unrooted trees, no node represents the ancestor and therefore no direction of evolution can be identified. A tree can be rooted by including an outgroup such as the most recent common ancestor. We rooted the tree by including one *Aspergillus oerlinghausenensis* and four *Aspergillus fischeri* genomes (Figure 2f, cluster 1). In three WGS studies of *A. fumigatus* ^25,27,32^ a genetically more distant cluster of *A. fumigatus* isolates has been identified (Figure 2f, cluster 2). To test if this cluster is monophyletic, Celia-Sanchez *et al.* sequenced eight housekeeping genes.^25^ For all cases the tree topology was consistent with polyphyly indicating that it does not have one common ancestor and therefore this cluster cannot be assigned as a clade. Triazole resistance is found throughout cluster 3 and 4 (Figure 2f), with triazole resistant *cyp*51A TR_34_ and TR_46_ haplotypes mainly in cluster 3. However, as triazole-sensitive *cyp*51A wild-type haplotypes are also found in this cluster, this cluster cannot be defined by TR triazole resistance only. Because the neighbor joining tree analysis is based on mutations and cannot consider recombination it forces the data to be shown in bifurcated branches. However, since *A. fumigatus* is a sexually recombining fungus, this branching is not a complete representation, and a phylogenetic network analysis should rather be used instead (Figure 2a-e).^24^

### Population structure associated with antifungal resistance

To quantify the genetic differentiation between subdivisions, we used F_ST_ or fixation index across windows of 10 kb. This compares the genetic structure of defined populations and patterns of evolutionary processes that shape existing genetic diversity. Comparing clinical versus environmental isolates (Figure 3a), no peaks (F_ST_ > 0·3) were observed, indicating the absence of differentiation between these two groups, confirming the previous results of Rhodes *et al.*^13^ However, when comparing triazole-susceptible to triazole-resistant isolates, roughly 36 significant genomic regions (average width 18,465 bp, min 4,999-max 77,499 bp) were observed (Supplemental Materials, Table S2). Although the *cyp*51A gene was not found in a differentiated region, manual inspection revealed the value of the TR_34_ variant position to be 0·40 and for TR_46_ 0·27, indicating differentiation of this locus between azole resistant and sensitive isolates, although not the nearby region. As our isolates were selected based on azole resistance phenotype, and *cyp*51A haplotype, we assumed that the *cyp*51A variants were causing the observed resistance. Although the general causality of this mechanism of resistance to triazoles has been confirmed by mutant analysis^33^, sexual crossing was carried out as a proof of concept (Supplemental Materials Table S3). The results of the crosses were consistent with triazole resistance being determined solely by *cyp*51A variants in nine of the ten isolates tested. Previously identified variants correlated with resistance to triazoles, such as HMG-CoA reductase-encoding gene, *hmg*1^S269F, S305P, G307D,^ were not observed in the Dutch *A. fumigatus* isolates. Therefore, although *hmg*1 or *hmg*2 mutations can confer resistance to triazoles, they are not genetically linked to the *cyp*51A gene TR_34_ or TR_46_ triazole resistance mechanisms in the Dutch population of *A. fumigatus* in this study.

**Figure 3.**
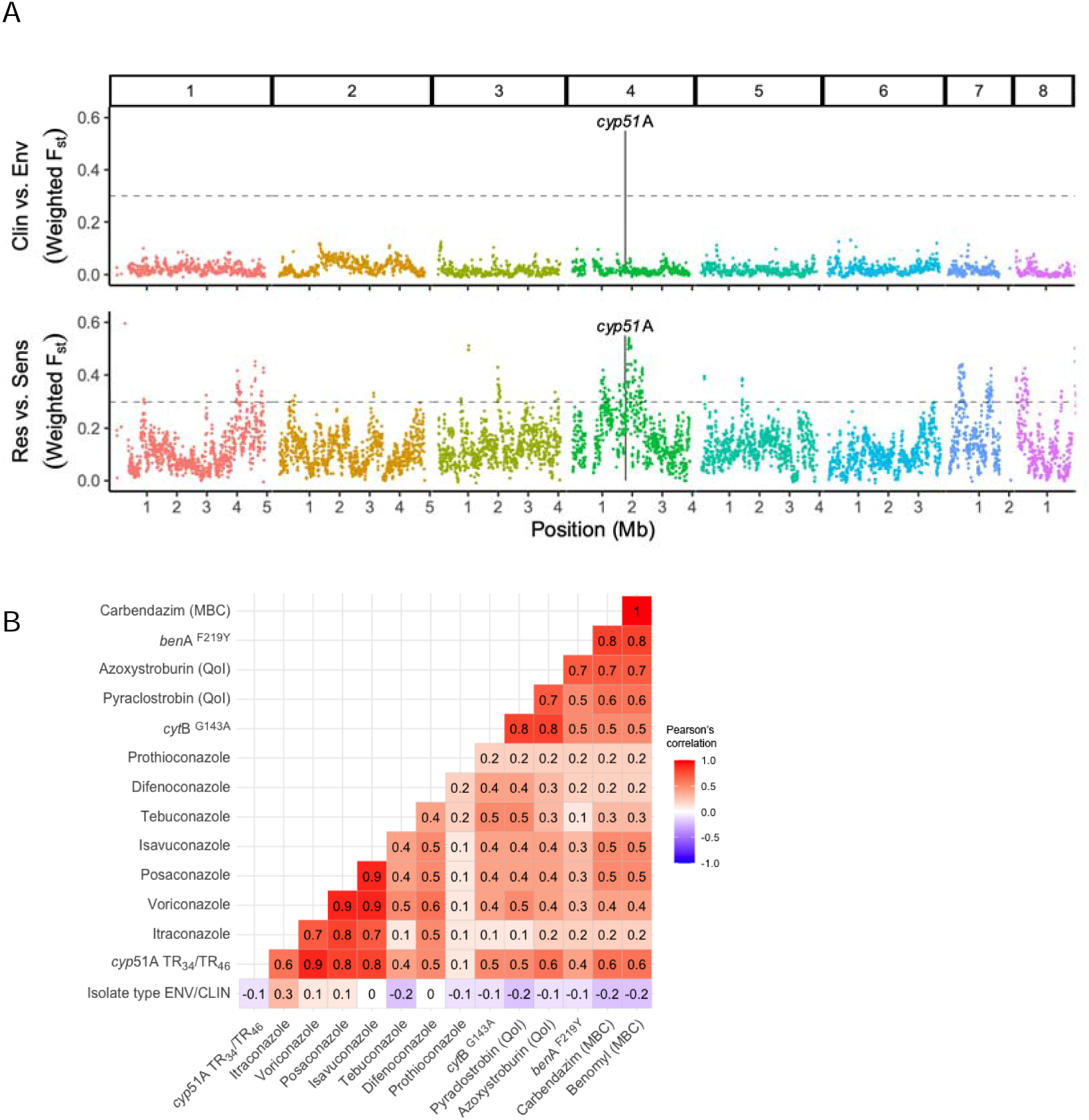
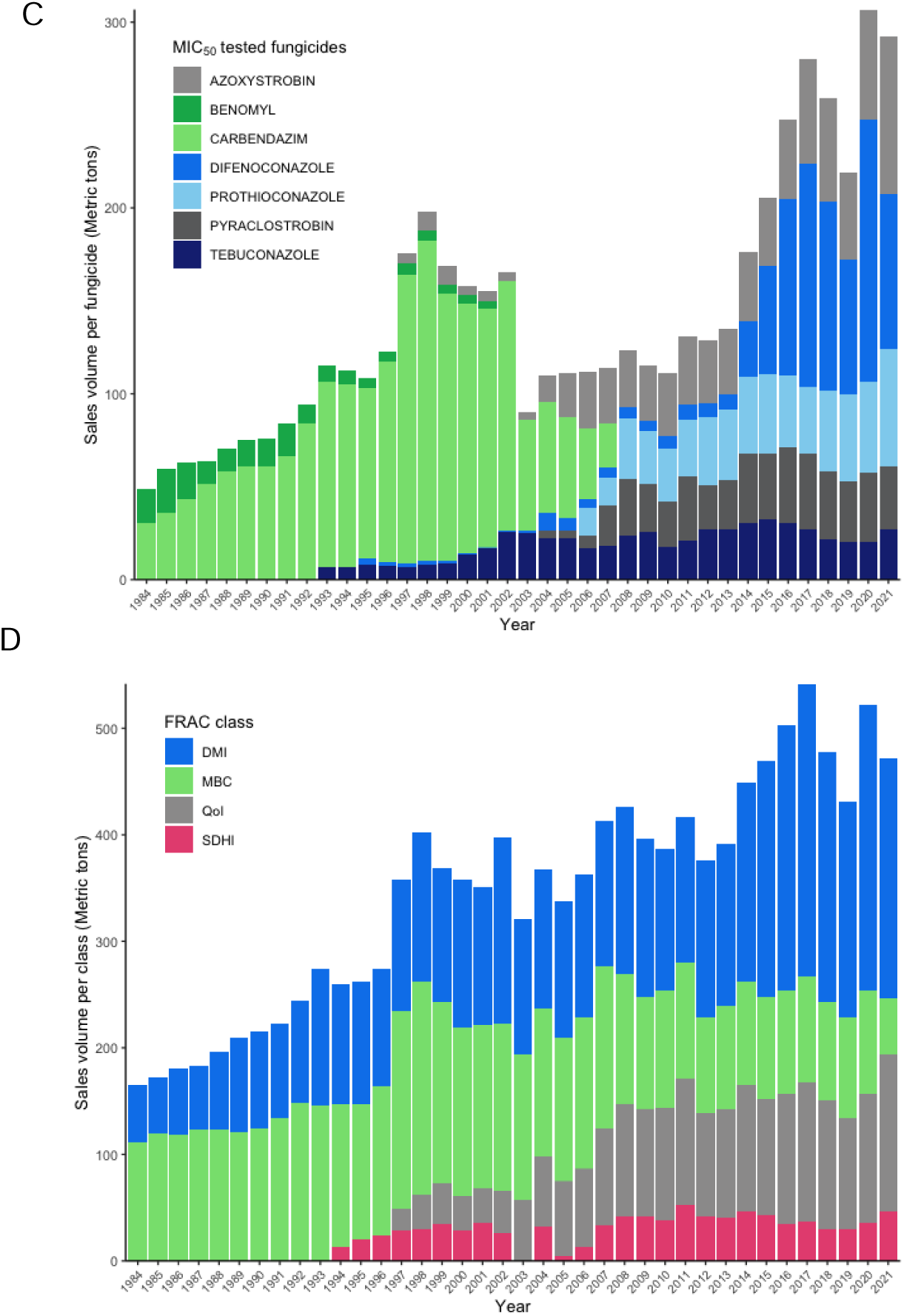
Correlation with multi-fungicide-resistance in *A. fumigatus*. a, F_ST_ analysis of 157 Dutch *A. fumigatus* isolates measuring the variance between different populations, split per chromosome (1-8). Top panel shows the comparison between environmental and clinical isolates with no peaks > 0·3, while the bottom panel shows triazole-resistant versus triazole-sensitive isolates with 36 significant regions with peaks > 0·3 (Supplemental Materials Table S2). b, Correlation plot of genotypic inferred resistance (TR_34_/TR_46_ *cyp*51A, *cyt*B^G143A^, *ben*A^F219Y^) and phenotypic resistance (MICs) and isolate type (environmental or clinical). As expected, a strong association is detected between medical triazoles phenotypes and *cyp*51A haplotypes, and the QoI and MBC resistance phenotype correlates with *cyt*B^G143A^, *ben*A^F219Y^ genotype. Additionally, a strong association is observed between triazoles and QoI and MBC genotypes and phenotypes. c, Sales volume in metric tons of the MIC tested fungicides in this study in the Netherlands between 1984 and 2021. In blue DMIs difenoconazole, prothioconazole and tebuconazole, in grey the QoIs azoxystrobin and pyraclostrobin, in green MBCs benomyl and carbendazim. d, Total sales volume in metric tons of fungicides per class of DMI (blue), QoI (grey), SDHI (pink) and MBC (green) in the Netherlands between 1984 and 2021 (data obtained from Dutch Ministry of Agriculture, Nature and Food quality).

A total of 36 differentiated regions (F_ST_ >0·3) were identified in the F_ST_ analysis when comparing phenotypically triazole-susceptible and triazole-resistant (Supplemental Materials Table S2). In 13 out of the 36 regions (putative) major facilitator superfamily (MFS) transporters and in three out of the 36 regions ATP-binding cassette (ABC) multidrug transporters were identified (Table S2 of Supplemental Materials). It is possible that these genes are also involved in resistance to other fungicide classes, and therefore, we inspected variants known to confer target-site fungicide resistance for quinone outside inhibitors (QoI), *cyt*B^G143A^, and methyl benzimidazole carbamate (MBC), *ben*A^F219Y^.^20,32,34^ In addition to genotyping all genomes (Figure 2f), we confirmed these resistances by phenotypic susceptibility testing of minimal inhibitory concentration (MIC) on a subset of 46 isolates. We then assessed correlations between genotypic resistance (*cyp*51A, *cyt*B, *ben*A) and phenotypic resistance (MIC_50_, Supplemental material Table S4) (Figure 3b). Similar to the PCA and F_ST_ analyses, there is no correlation between the source of an isolate (environmental/clinical) and its geno- or phenotype. The strongest associations were observed between medical triazole resistance phenotypes and the *cyp*51A TR_34_/TR_46_ alleles, while no associations were observed for prothioconazole. Other strong associations were observed between the QoI compounds and *cyt*B^G143A^, and MBC compounds and the *ben*A^F219Y^ resistance allele. Interestingly, there is also a strong association observed between triazoles and QoI and MBC geno- and phenotypes. When an isolate is phenotypically resistant (MIC) and carrying a haplotype resistant to triazoles (*cyp*51A) it is more likely to also be resistant to QoIs and MBCs. (Table S4). Looking at all the Dutch genomes of this study (n=157), 31% of the triazole-resistant *A. fumigatus* isolates with a TR_34_ or TR_46_ *cyp*51A haplotype have both the *cyt*B^G143A^ and *ben*A^F219Y^ variants (9% with either one or the other) that give rise to resistance to the QoI compounds pyraclostrobin and azoxystrobin and the MBC compounds carbendazim and benomyl. To study the use of these particular fungicide compounds in the Netherlands, we obtained the fungicide sales data in the Netherlands for the past 30 years (1984 – 2009 data obtained from archives of Dutch Ministry of Agriculture, Nature and Food quality, 2010 – 2021 data obtained from www.rijksoverheid.nl). Of the MIC tested fungicides in this study we show its specific sales volume per fungicide from 1984 till 2021 in the Netherlands in Figure 2c. Over the years individual sales of fungicides fluctuate and some fungicides have been discontinued such as benomyl and carbendazim of the MBC class, although we still detect phenotypic resistance to these compounds in *A. fumigatus*. Looking at the yearly sales data per whole class of these fungicides in the Netherlands, according to the Fungicide Resistance Action Committee (FRAC) classification (Figure 2d), the MBC class has still been sold in the past decade and could therefore explain the presence of the resistance allele in the *A. fumigatus* population in the Netherlands. The SDHI class remains stable at a relatively low level while the DMI and QoI are increasing in sales volume overall. We detected no phenotypic resistance to the dihydroorotate dehydrogenase (DHODH) inhibitor olorofim (Supplemental Materials Table S4).^35^

## Discussion

This study strengthens earlier findings of clonal groups of environmental and clinical isolates and strongly suggests the transmission of the triazole-resistant TR_34_ and TR_46_ *cyp*51A haplotypes from the environment to patients.^25,29^ Here, we show for the first time clonal groups in which clinical isolates that caused probable aspergillus disease are nearly identical to environmental isolates. Although we were able to recover clonal groups, a primary goal of genomic epidemiology, geography and timeline appear to be insufficient to explain direct transmission routes of *A. fumigatus*. In 1998 a study using DNA fingerprinting of more than 700 French *A. fumigatus* isolates concluded that there was not a clear correlation between geography and isolate genotype. However, they recovered identical isolates shared between the environment and patients.^36^ This study was the first to our knowledge to show identical types across larger distances, reproduced decades later by large scale use of microsatellites.^14,37,38^ These studies agree that recovery of clonal isolates is insufficient to identify a patient’s source of infection – since identical microsatellite profiles can be found in multiple countries. In the current study, using WGS on Dutch *A. fumigatus* isolates and including publicly available genomes too, a total of 205 clonal groups were detected across the globe. We found that for three of the four Dutch clonal groups, additional clones were identified in other European samplings. It is therefore incorrect to singly target a particular Dutch agricultural site as the source, simply because the sample was taken in the Netherlands. In the absence of widespread routine genomic monitoring, we lack an overview of the potential sources and ecology of this fungus, specifically at the local level necessary to explain finer scale differences. Unfortunately, this dispersal [absence of local groups] implies that genomic epidemiology alone is of limited value for determining the origin and transmission route of individual *A. fumigatus* infections.

Population genomic analysis supports a highly interconnected population, with some level of population structure in *A. fumigatus*, but without evidence of clades.^13,21,22,24,25^ The correlation of triazole-resistance and multi-fungicide-resistance in individual isolates could be driven by fungicide exposure, though other agricultural practices may also have been a significant contributor. The high use of fungicides to enable the transition to intensive agriculture in the past century appears to have created novel agricultural environmental niches with strong selection pressure for fungicide resistance.^39^ Our results confirm, as previously shown, the presence of additional, non-triazole, resistance mechanisms, such as QoI and MBC resistance within *A. fumigatus*, an off-target fungus for these agricultural fungicides.^20,32,34,40^ The association between other fungicide resistance alleles and triazole resistance in *A. fumigatus* raises concerns about the broader impact of fungicide usage on the emergence and persistence of antifungal resistance in environmental settings. Due to this genetic correlation of resistances, discontinuing the use of triazoles in the environment alone may not have a significant impact on the reduction of TR_34_ and TR_46_ *cyp*51A haplotypes, so long as these other classes of fungicides are still applied. The persistence of resistance highlights a potential barrier for public policies. Furthermore, the simultaneous development of new modes of action for medical and agricultural applications underscores the continued risk for cross resistance.^35^ Policymakers, researchers, and practitioners should recognize that the consequences of fungicide resistance extend beyond their immediate applications and have long-term implications for human health.

## Contributors

Conceptualization: ES, BZ and PV

Data curation: ES, YN and BA

Formal analysis: ES, BCS, YN, LW and BA

Funding acquisition: ES, BZ, PV, MM, and MB

Investigation: ES, BCS, YN, JZ, HK, MKT, AGL, LW and BA

Methodology: ES, JZ, BA and PV

Project administration: ES

Resources: ES, KvD and PV

Software: BA

Supervision: ES, MM, and MB Visualization: ES, BA

Writing - original draft: ES, BA, PV

Writing - review and editing: ES, JZ, HK, BZ, AGL, KvD, LW, MB, MM, BA, PV

## Data sharing statement

Genomes have been deposited under project PRJEB73793 at ENA. All bioinformatical scripts and analysis are available at https://github.com/fungalsnelderslab/2024_Dutch_fumigatus. The Dutch *A. fumigatus* isolates of this study are available upon request for environmental isolates, for clinical isolates.

## Acknowledgments

Paul van der Wielen from KWR for providing *A. fumigatus* isolates from water. M.M., M.T.B. and B.N. C-S. were supported by funding from the Centers for Disease Control and Prevention (CDC; contract 0HCVLD13-2018-27470 to M.M. and M.T.B.) and United States Department of Agriculture, National Institute of Food and Agriculture (USDA NIFA AFRI grant 2019-67017-29113 to M.T.B. and M.M.). B.N.C-S. was also supported by the National Science Foundation under Grant No. DGE-1545433. We would like to thank Prof. Matthew Fisher and Dr. Johanna Rhodes for input on a previous version of this manuscript.

## References

1 Verweij PE, Brüggemann RJM, Azoulay E, et al. Taskforce report on the diagnosis and clinical management of COVID-19 associated pulmonary aspergillosis. Intensive Care Med 2021; 47: 819–34.

2 Verweij PE, Rijnders BJA, Brüggemann RJM, et al. Review of influenza-associated pulmonary aspergillosis in ICU patients and proposal for a case definition: an expert opinion. Intensive Care Med 2020; 46: 1524–35.

3 Latgé J-P, Chamilos G. Aspergillus fumigatus and Aspergillosis in 2019. Clin Microbiol Rev 2019; 33: e00140–18.

4 Herbrecht R, Denning DW, Patterson TF, et al. Voriconazole versus Amphotericin B for Primary Therapy of Invasive Aspergillosis. N Engl J Med 2002; 347: 408–15.

5 Snelders E, van der Lee HAL, Kuijpers J, et al. Emergence of Azole Resistance in Aspergillus fumigatus and Spread of a Single Resistance Mechanism. PLoS Med 2008; 5: e219.

6 Snelders E, Camps SMT, Karawajczyk A, et al. Triazole Fungicides Can Induce Cross-Resistance to Medical Triazoles in Aspergillus fumigatus. PLoS ONE 2012; 7: e31801.

7 Schoustra SE, Debets AJM, Rijs AJMM, et al. Environmental Hotspots for Azole Resistance Selection of Aspergillus fumigatus , the Netherlands. Emerg Infect Dis 2019; 25: 1347–53.

8 van der Linden JWM, Camps SMT, Kampinga GA, et al. Aspergillosis due to Voriconazole Highly Resistant Aspergillus fumigatus and Recovery of Genetically Related Resistant Isolates From Domiciles. Clin Infect Dis 2013; 57: 513–20.

9 van Ingen J, van der Lee HA, Rijs TAJ, et al. Azole, polyene and echinocandin MIC distributions for wild-type, TR34/L98H and TR46/Y121F/T289A Aspergillus fumigatus isolates in the Netherlands. J Antimicrob Chemother 2015; 70: 178–81.

10 Kolwijck E, van der Hoeven H, de Sévaux RGL, et al. Voriconazole-Susceptible and Voriconazole-Resistant Aspergillus fumigatus Coinfection. Am J Respir Crit Care Med 2016; 193: 927–9.

11 Lestrade PP, Bentvelsen RG, Schauwvlieghe AFAD, et al. Voriconazole Resistance and Mortality in Invasive Aspergillosis: A Multicenter Retrospective Cohort Study. Clin Infect Dis 2019; 68: 1463–71.

12 Barber AE, Sae-Ong T, Kang K, et al. Aspergillus fumigatus pan-genome analysis identifies genetic variants associated with human infection. Nat Microbiol 2021; 6: 1526–36.

13 Rhodes J, Abdolrasouli A, Dunne K, et al. Population genomics confirms acquisition of drug-resistant Aspergillus fumigatus infection by humans from the environment. Nat Microbiol 2022; 7: 663–74.

14 Rocchi S, Sewell TR, Valot B, et al. Molecular Epidemiology of Azole-Resistant Aspergillus fumigatus in France Shows Patient and Healthcare Links to Environmentally Occurring Genotypes. Front Cell Infect Microbiol 2021; 11: 729476.

15 Lavergne R-A, Chouaki T, Hagen F, et al. Home Environment as a Source of Life-Threatening Azole-Resistant Aspergillus fumigatus in Immunocompromised Patients: Table 1. Clin Infect Dis 2017; 64: 76–8.

16 Zhang J, Lopez Jimenez L, Snelders E, et al. Dynamics of Aspergillus fumigatus in Azole Fungicide-Containing Plant Waste in the Netherlands (2016–2017). Appl Environ Microbiol 2021; 87. DOI:10.1128/AEM.02295-20.

17 Zhang J, Debets AJM, Verweij PE, Schoustra SE. Selective Flamingo Medium for the Isolation of Aspergillus fumigatus. Microorganisms 2021; 9: 1155.

18 Buil JB, van der Lee H a. L, Rijs AJMM, et al. Single-Center Evaluation of an Agar-Based Screening for Azole Resistance in Aspergillus fumigatus by Using VIPcheck. Antimicrob Agents Chemother 2017; 61: e01250–17.

19 Tang CM, Cohen J, Holden DW. An Aspergillus fumigatus alkaline protease mutant constructed by gene disruption is deficient in extracellular elastase activity. Mol Microbiol 1992; 6: 1663–71.

20 Kang SE, Sumabat LG, Melie T, Mangum B, Momany M, Brewer MT. Evidence for the agricultural origin of resistance to multiple antimicrobials in Aspergillus fumigatus , a fungal pathogen of humans. G3 GenesGenomesGenetics 2022; 12: jkab427.

21 Zhao S, Ge W, Watanabe A, Fortwendel JR, Gibbons JG. Genome-Wide Association for Itraconazole Sensitivity in Non-resistant Clinical Isolates of Aspergillus fumigatus. Front Fungal Biol 2021; 1: 617338.

22 Etienne KA, Berkow EL, Gade L, et al. Genomic Diversity of Azole-Resistant Aspergillus fumigatus in the United States. mBio 2021; 12: e01803–21.

23 Lofgren LA, Lorch JM, Cramer RA, et al. Avian-associated Aspergillus fumigatus displays broad phylogenetic distribution, no evidence for host specificity, and multiple genotypes within epizootic events. G3 GenesGenomesGenetics 2022; 12: jkac075.

24 Lofgren LA, Ross BS, Cramer RA, Stajich JE. The pan-genome of Aspergillus fumigatus provides a high-resolution view of its population structure revealing high levels of lineage-specific diversity driven by recombination. PLOS Biol 2022; 20: e3001890.

25 Celia-Sanchez B, Mangum B, Gómez Londoño L, et al. Pan-azole-and multi-fungicide-resistant Aspergillus fumigatus is widespread in the United States. Appl Environ Microbiol 2024; 90: e01782–23.

26 Barber AE, Riedel J, Sae-Ong T, et al. Effects of Agricultural Fungicide Use on Aspergillus fumigatus Abundance, Antifungal Susceptibility, and Population Structure. mBio 2020; 11: e02213–20.

27 Barber AE, Sae-Ong T, Kang K, et al. Aspergillus fumigatus pan-genome analysis identifies genetic variants associated with human infection. Nat Microbiol 2021; 6: 1526–36.

28 Zhang J, Verweij PE, Rijs AJMM, Debets AJM, Snelders E. Flower Bulb Waste Material is a Natural Niche for the Sexual Cycle in Aspergillus fumigatus. Front Cell Infect Microbiol 2022; 11: 785157.

29 Rhodes J, Abdolrasouli A, Dunne K, et al. Population genomics confirms acquisition of drug-resistant Aspergillus fumigatus infection by humans from the environment. Nat Microbiol 2022; 7: 663–74.

30 Kortenbosch HH, Van Leuven F, Van Den Heuvel C, Schoustra SS, Zwaan BJ, Snelders E. Catching more air: A method to spatially quantify aerial triazole resistance in Aspergillus fumigatus. 2022; published online Nov 4. DOI:10.1101/2022.11.03.515058.

31 Shelton JMG, Rhodes J, Uzzell CB, et al. Citizen science reveals landscape-scale exposures to multiazole-resistant Aspergillus fumigatus bioaerosols. Sci Adv 2023; 9: eadh8839.

32 Gonzalez-Jimenez I, Garcia-Rubio R, Monzon S, Lucio J, Cuesta I, Mellado E. Multiresistance to Nonazole Fungicides in Aspergillus fumigatus TR _34_ /L98H Azole-Resistant Isolates. Antimicrob Agents Chemother 2021; 65: e00642–21.

33 Snelders E, Karawajczyk A, Verhoeven RJA, et al. The structure–function relationship of the Aspergillus fumigatus cyp51A L98H conversion by site-directed mutagenesis: The mechanism of L98H azole resistance. Fungal Genet Biol 2011; 48: 1062–70.

34 Fraaije B, Atkins S, Hanley S, Macdonald A, Lucas J. The Multi-Fungicide Resistance Status of Aspergillus fumigatus Populations in Arable Soils and the Wider European Environment. Front Microbiol 2020; 11: 599233.

35 Verweij PE, Arendrup MC, Alastruey-Izquierdo A, et al. Dual use of antifungals in medicine and agriculture: How do we help prevent resistance developing in human pathogens? Drug Resist Updat 2022; 65: 100885.

36 Chazalet V, Debeaupuis J-P, Sarfati J, et al. Molecular Typing of Environmental and Patient Isolates of Aspergillus fumigatus from Various Hospital Settings. J Clin Microbiol 1998; 36: 1494–500.

37 Sewell TR, Zhu J, Rhodes J, et al. Nonrandom Distribution of Azole Resistance across the Global Population of Aspergillus fumigatus. mBio 2019; 10: e00392–19.

38 Fraaije BA, Atkins SL, Santos RF, Hanley SJ, West JS, Lucas JA. Epidemiological Studies of Pan-Azole Resistant Aspergillus fumigatus Populations Sampled during Tulip Cultivation Show Clonal Expansion with Acquisition of Multi-Fungicide Resistance as Potential Driver. Microorganisms 2021; 9: 2379.

39 Fisher MC, Alastruey-Izquierdo A, Berman J, et al. Tackling the emerging threat of antifungal resistance to human health. Nat Rev Microbiol 2022; 20: 557–71.

40 Schürch S, Gindro K, Schnee S, et al. Occurrence of Aspergillus fumigatus azole resistance in soils from Switzerland. Med Mycol 2023; 61: myad110.

