## Supplemental methods & materials for "Widely dispersed clonal expansion of multi-fungicide-resistant *Aspergillus fumigatus* limits genomic epidemiology prospects"

**Sample selection**

After initial sample selection, to match 1:1 clinical and environmental isolates with respect to *cyp*51A genotype and triazole susceptibility, it was necessary to extend the year of culture of clinical isolates up to 2020 to include an additional 43 isolates to reach a total of 85 clinical isolates (Supplemental Materials Figure S1). An overview of the basic sequencing statistics of the 174 samples and excluded isolates can be found in Supplemental Methods. One sample was removed due to low coverage (<10X coverage). By using a reference-guided assembly (Af293) and multiple pairwise comparison method, we found on average between 5-1000 unique single nucleotide variants (SNV). In one case 49,456 unique SNVs were observed and this isolate was removed as preliminary analyses showed it was part of a described divergent population.(1) Another 15 whole genome sequenced isolates were removed due to high levels of heterozygosity, potentially indicating mixed samples, as on average 20% of the total sequence was called heterozygous, but for 15 isolates this was higher than 35% and these isolates were removed for all further analyses. Thus, 157 of the initial 174 whole genome sequences were retained and used in all analyses and metadata regarding the distribution of the remaining 157 clinical and environmental samples are found in Supplementary Excel file 1.

To this Dutch dataset, publicly available samples were added, and reads were aligned, mapped, and variants were quality filtered as above. Triazole resistance phenotypes for these global samples were obtained from the respective publication, where available.(1–9)

**Bioinformatic analysis**

To assess the genetic diversity, Illumina paired-end reads were mapped to the Af293 reference genome assembly (ASM265v1), to which we had added the mitochondrial genome (JQ346808.1). Reads were mapped using bwa-mem2 v2.0pre2 (10), and duplicates marked with samtools v1.10 (11). Variants were identified using GATK v4.2.6.1 (12) for each isolate with the GVCF mode, and then combined prior to joint genotyping. Following joint genotyping, variants were separated into SNP and INDEL categories and SNPs filtered based on the command “QD < 2.0 || MQ < 40.0 || FS > 60.0 || MQRankdSum < -12.5 || ReadPosRankSum < -8.0 || AN < 300” while indelss were filtered on “QD < 2.0 || FS > 200.0 || ReadPosRankSum < -20.0 || AN < 300 “. These SNPs and indels were merged for the final filtered variant dataset.

For quality control purposes, we removed 18 samples (7 environmental; 11 clinical) where more than 10% of the variants were heterozygous, which likely indicates mixed cultures. For the remaining 156 samples the average genome-wide sequence coverage was 48.4X. The filtered variants resulted in a per-sample average of 49,422 SNPs and 5,325 Indels compared to the Af293 reference.

The filtered variants were clustered based on genomic Principal Components using plink v1.9 (13), and visualized with R v4.2.1 (14) using the tidyverse package v1.3.2 (15). To visualize the relatedness between isolates within this recombining species, a phylogenetic network was constructed using splitstree v4.19.0 (16), after converting the VCF to a phylip format with a script from <https://github.com/edgardomortiz/vcf2phylip>.(17) To explore the effect of biased sampling on phylogenetic network visualization, we subsetted the global variant dataset based on samples with published azole resistance, picking approximately 350 samples for each subsetted ratio. This subsetted variant dataset was converted to a phylip file and used for splistree as above.

To assess the relationships between mutations and population structure, although this is a sexually reproducing species, we used IQ-TREE to produce a neighbor-joining tree (-fast option) using the Jukes-Cantor substitution model.(18) The tree was visualized with associated phenotypic/genetic data using ggtree.(19)

As a measure of population differentiation within the Dutch *A. fumigatus* population, *F_ST_* was calculated in sliding windows of 10 kb using vcftools v0.1.16 (20). The samples were split based on two conditions, clinical versus environmental source, or azole resistant or azole sensitive. For the *F_ST_* calculation, the azole resistance/sensitivity was classified based on resistance to at least one of voriconazole, itraconazole, or posaconazole on the VIP check plates (21). Results of the *F_ST_* calculation were plotting using R v4.2.1.(14)

To visualize all globally clonal or highly identical isolates, UpSetR was used (22). The UpSet technique visualized groupings of clonal groups of *A. fumigatus* isolates from The Netherlands, UK, USA and Japan visualized with R v4.2.1 (14) using the UpSetR package 1.4.0 (14,23).

***Cyp*51A segregation sexual crossing experiments**

A large set of azole sensitive *A. fumigatus* isolates were one by one crossed with azole resistant isolates (this study) from opposing mating type on oatmeal agar (Difco, Oatmeal Agar 72 g/L). After incubation of six to ten eight weeks, cleistothecia were carefully removed and suspended in 0.05% saline-tween and heated in a heat block at 70°C for 1 hour to kill off any remaining conidiospores. Not all crosses resulted in successful sexual cycles but among the successful crosses, we selected twelve pairings between an azole sensitive parent and a resistant parent, six with the TR_34_/L98H and six with the TR_46_/Y121F/T289A *cyp*51A haplotypes. All phenotypically azole sensitive parents had wild type *cyp*51A haplotypes. Per selected cross, the ascospores were plated out on MEA medium and cultured till the colonies showed early signs of minimal sporulation. For most of the offspring, this was around 40 hours of culturing at 37°C, or 48 hours of culturing at 30°C. A minimum of 32 colonies were pre-cultured per cross and transferred to two sets of Petri dishes containing 50% MEA, one set being azole negative, and the other set containing 8mg/L itraconazole. Each plate was marked with a 4x4 numbered square grid; the points on the grid were spaced 15mm apart. For phenotyping, each of the 32 colonies was transferred to a correspondingly numbered dot on both an azole-negative and azole-positive plate. Phenotyping plates were cultured for 48 hours at 37°C. After which growth on the azole-positive plates was used to score resistance. Growth with a diameter of >1 mm at the inoculation point was scored as resistant, <1 mm was scored as sensitive. From a random subset of 16 out of the 32 offspring, conidiospores were harvested from the control colonies on the azole negative plates using cotton swabs wetted with PBS-tween. The spores were suspended in 50 µl of PBS-tween in a PCR 96-well plate. Subsequently, DNA was extracted from the spore suspension using the heatshock DNA extraction protocol as described in Basic protocol 2 in Fraczek *et al.* (24), except that the suspensions were cooled in a -20°C freezer for an hour rather than in a -80°C freezer for 20 minutes after the heat shock. The promotor region of the *cyp*51A gene was amplified as described previously (25). For genotyping the offspring as wildtype (WT) or tandem repeat (TR), 3 µL PCR product of each sample was loaded onto a 1% agarose gel. Gels were run for 50 minutes at 100V. The 34-or 46 bp difference in fragment size between the WT or TR fragments was visually scored. A Chi-square test was performed to test for a significant deviation from a 1:1 phenotype ratio between the number of sensitive and resistant offspring. Statistical analysis and generation of summary tables were performed in R version 4.1.3 (14). The following R packages: reshape2 1.4.4 (15) and janitor 2.1.0 (26), were used for the analysis.

**Microdilution antifungal susceptibility testing**

From the total 157 isolates of this study, a selection of 46 isolates were tested for phenotypic susceptibility to medical triazoles, agricultural triazoles; demethylase inhibitors (DMI), quinone outside inhibitors (QoI) and methyl benzimidazole 2-yl carbamate (MBC) fungicides. 23 clinical (Clin) and 23 environmental (Env) isolates of which; 8 Clin and 8 Env *cyp*51A wild type *A. fumigatus* isolates, 1 Clin G43R, 11 Clin and 11 Env triazole resistant TR_34_/L98H, 3 Clin and 4 Env TR_46_/Y121F/T289A isolates (Supplemental Materials Table S4). The *in vitro* activity of itraconazole, voriconazole, posaconazole, isavuconazole, olorofim, tebuconazole, difenoconazole, prothioconazole, pyraclostrobin, azoxystrobin, carbendazim, benomyl (Sigma Aldrich) was tested. The minimal inhibiting concentration (MIC) was determined using a microbroth dilution format according to the EUCAST reference method E.DEF 9.4;
[www.eucast.org/astoffungi/methodsinantifungalsusceptibilitytesting/ast_of_moulds](http://www.eucast.org/astoffungi/methodsinantifungalsusceptibilitytesting/ast_of_moulds). MIC was determined at 100% growth inhibition (MIC_100_) for medical antifungal compounds and 50% for agricultural antifungal compounds (MIC_50_).

**Correlation plot**

A correlation plot of genotypic resistance (*cyp*51A, *cyt*B, *ben*A) and phenotypic resistance (MIC of itraconazole, voriconazole, posaconazole, isavuconazole, olorofim, tebuconazole, difenoconazole, prothioconazole, pyraclostrobin, azoxystrobin, carbendazim, benomyl) of a subselection of 46 *A. fumigatus* isolates (see Microdilution antifungal susceptibility testing). MIC data was log_2_ transformed and a pairwise comparison was done by using corrplot version 0.92 (27). Results of this pairwise comparison were plotted using ggcorrplot (28) version 3.4.4 in R v4.2.1 (14).

**Fungicides sales data**

A sales data set of all active compounds listed as fungicides in the Netherlands under “Meerjarenplan gewasbescherming” (MJP-G) from 1984 to 2021 were used to plot agricultural triazoles, methyl benzimidazol-2-yl carbamate (MBC) and Quinone outside Inhibitor (QoI) sold in the Netherlands (1984 – 2009 data obtained archives of the Dutch Ministry of Agriculture, Nature and Food, 2010 – 2021 data obtained from www.rijksoverheid.nl). Only data of fungicides that were MIC_50_ tested in this study were plotted in figure 3c, for triazoles (DMI); difenoconazole, tebuconazole, MBCs; benomyl and carbendazim, and QoIs azoxystrobin and pyroclostrobin. In figure 3d all fungicides sold in the Netherlands according to this dataset were plotted for the classes DMI, MBC, QoI and SHDI. Sales was visualized in a stacked bar plot by year using ggplot2 (28) version 3.4.4 in R v4.2.1 (14).

**Supplemental Materials**

**Widely dispersed clonal expansion of multi fungicide resistant *Aspergillus fumigatus* limits genomic epidemiology prospects**

Eveline Snelders^1^, Brandi Nicole Celia-Sanchez^2^, Ymke Nederlof^1^, Jianhua Zhang^1^, Hylke Kortenbosch^1^, Marlou Tehupeiory-Kooreman^3^, Li Wang^4^, Karin van Dijk^5^, Marin Talbot Brewer^4^, Michelle Momany^2^, Bas J Zwaan^1^, Ben Auxier^1^, Paul E. Verweij^3^

Supplementary datafile:

Baseclear_metadata

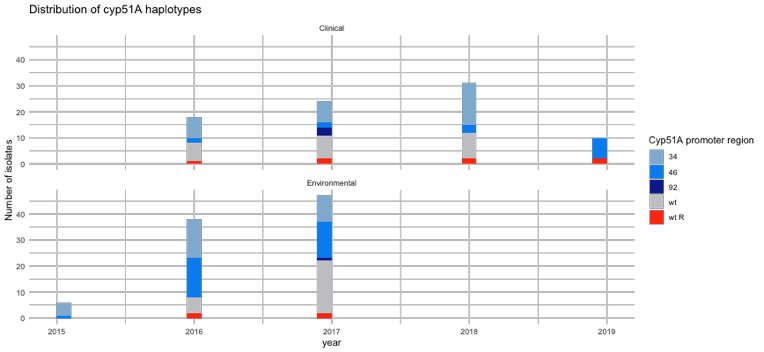

**Figure S1. Distribution of sample selection**

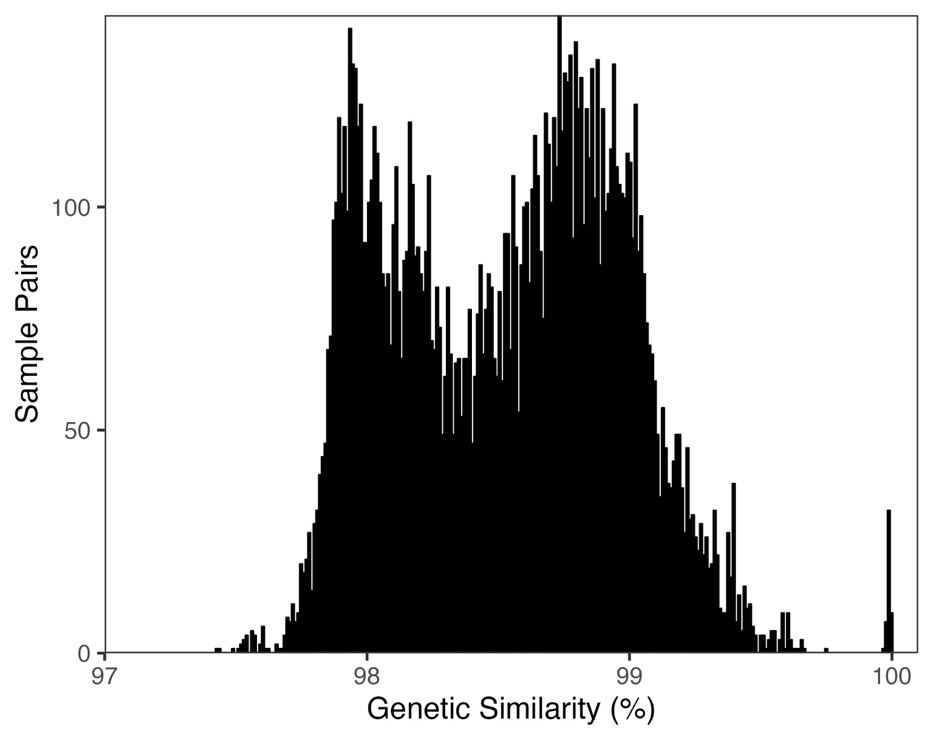

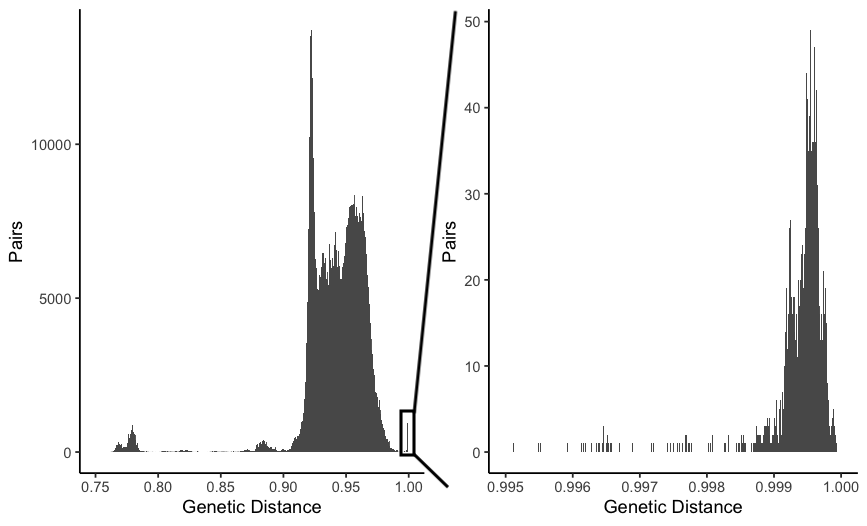

**Figure S2.** Histogram pairwise comparison between all 157 *A. fumigatus* isolates (top panel) and all 1231 global isolates (bottom panel) on the left bottom all global comparisons and on the right a zoomed in image of all global clonal groups.

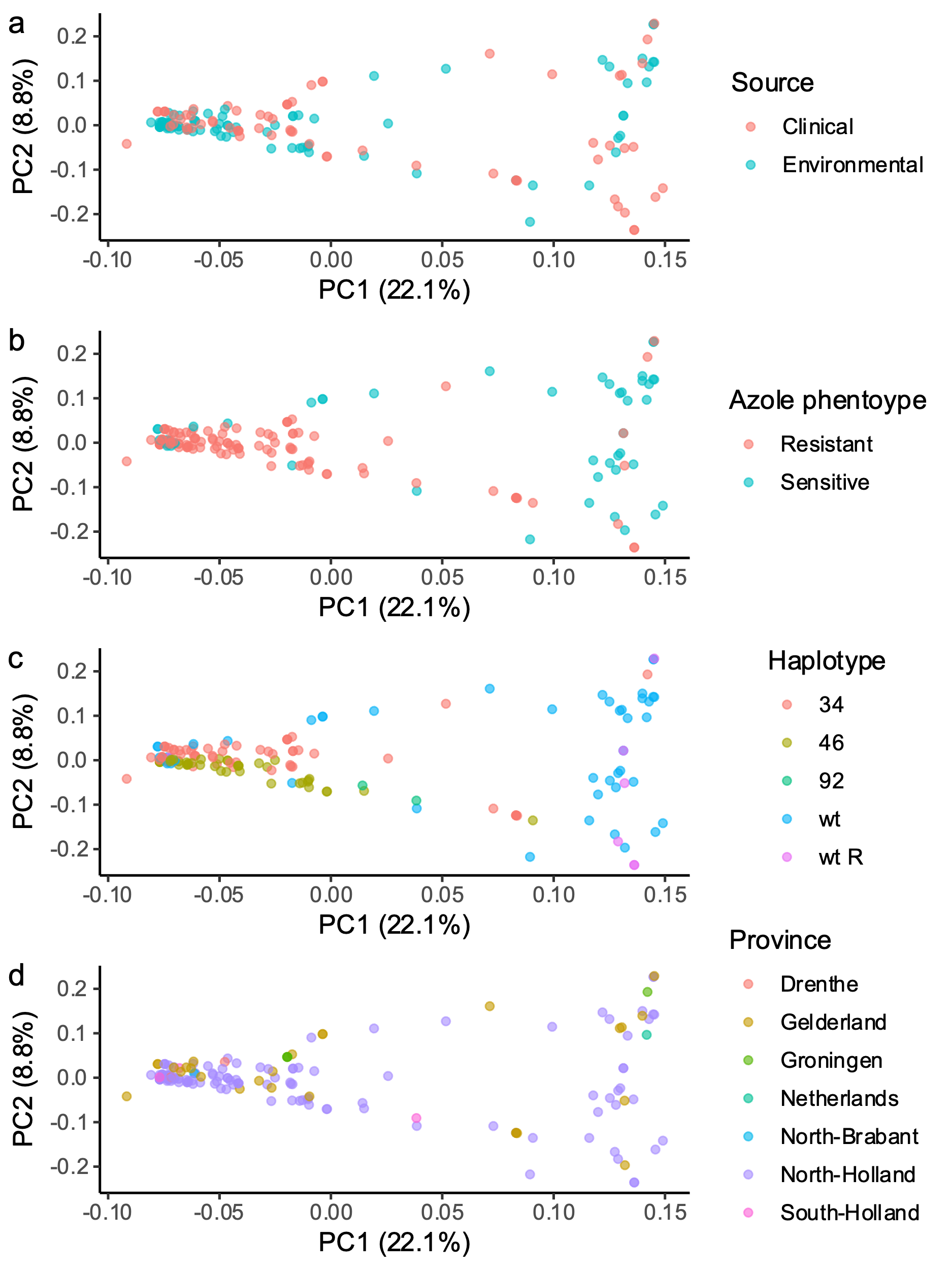

**Figure S3.** Principle component analysis (PCA) of 159 Dutch *A. fumigatus* genomes of azole resistant versus sensitive isolates (a), clinical versus environmental isolates (b) on *cyp*51A haplotype, and geographic location of isolates in the Netherlands (d)

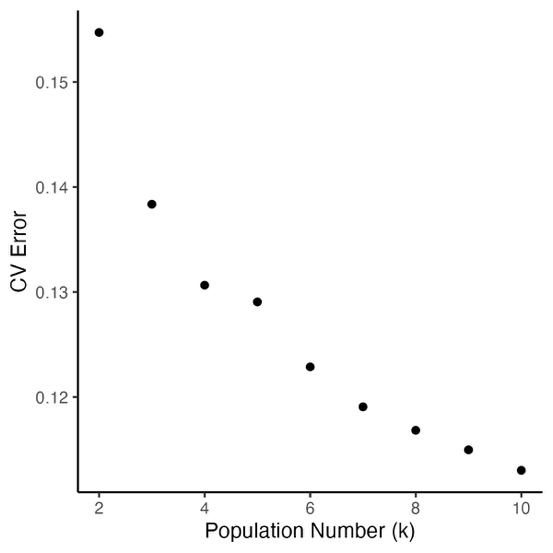

**Figure S4. ADMIXTURE analysis of the global dataset using different values of k.**

**Table S1.** Azole phenotype, strain identity, site and year of isolation, and *Aspergillus* disease classification.

| **Year** | **2016** | | **2017** | | **2018** | | **2019** | |  |
| --- | --- | --- | --- | --- | --- | --- | --- | --- | --- |
| Site / category | Strain ID | Disease classification | Strain ID | Disease classification | Strain ID | Disease classification | Strain ID | Disease classification | Total |
| **WT** |  |  |  |  |  |  |  |  |  |
| AmsterdamUMC | 513 | Probable IPA | 1464 | ABPA | 162 | IA |  |  |  |
|  | 1545 | Probable IPA | 1730 | Non-invasive | 980 | IA |  |  |  |
|  | 2406 | CNS IA | 1976 | Proven IA | 1267 | Fungal mastoiditis |  |  |  |
|  |  |  | 2108 | Probable IA | 1533 | ABPA |  |  |  |
|  |  |  | 2109 | Probable IA | 1945 | IA |  |  |  |
|  |  |  |  |  | 2288 | Aspergilloma |  |  |  |
| subtotal | 3 |  | 5 |  | 6 |  |  |  | 14 |
| Radboudumc | V212-54 | Proven IA | V234-13 | Proven IA | V250-64 | IA |  |  |  |
|  | V208-52 | Proven IA | V232-75 | Putative IA | V254-55 | Probable IA |  |  |  |
|  | V207-11 | Probable IA | V231-63 | Probable IA | V254-50 | Putative IA |  |  |  |
|  | V197-79 | Proven IAPA | V228-47 | Proven IA | V240-10 | Probable IA |  |  |  |
| subtotal | 4 |  | 4 |  | 4 |  |  |  | 12 |
| **TR_34_/L98H** |  |  |  |  |  |  |  |  |  |
| AmsterdamUMC | V209-56 | Probable IA | V219-48 | Probable IA | V265-12 | Probable IA |  |  |  |
|  | V199-25 | No IA | V229-37 | Probable IA | V265-56 | Probable IA |  |  |  |
|  | V200-50 | No IA | V215-18 | Probable IA | V261-31 | No IA |  |  |  |
|  | V210-08 | No IA | V238-07 | Aspergilloma |  |  |  |  |  |
| subtotal | 4 |  | 4 |  | 3 |  |  |  | 11 |
| Radboudumc | V192-81 | Probable IA | V215-11 | Putative IA | V251-74 | Probable IA |  |  |  |
|  | V196-05 | Probable IA | V215-62 | Proven IA | V255-23 | Probable IA |  |  |  |
|  | V198-65 | Proven IA | V224-66 | Putative IA | V257-33 | Probable IA |  |  |  |
|  | V200-72 | Colonization | V233-43 | Colonization | V261-45 | Probable IA |  |  |  |
|  |  |  |  |  | V266-66 | Probable IA |  |  |  |
| subtotal | 4 |  | 4 |  | 5 |  |  |  | 13 |
| **TR_46_/Y121F/T289A** |  |  |  |  |  |  |  |  |  |
| AmsterdamUMC | V204-54 | No IA |  |  | V258-36 | Probable IA | V275-41 | IAPA |  |
|  |  |  |  |  | V252-43 | No IA | V289-01 | No IA |  |
|  |  |  |  |  | 261-15 | No IA | V294-36 | No IA |  |
|  |  |  |  |  |  |  | V297-39 | No IA |  |
|  |  |  |  |  |  |  | V306-43 | Probable IA |  |
|  |  |  |  |  |  |  | V306-67 |  |  |
| subtotal | 1 |  | 0 |  | 3 |  | 6 |  | 10 |
| Radboudumc | V213-14 | Colonization | V226-13 | Proven IA |  |  | V282-59 | Putative IAPA |  |
|  |  |  | V226-28 | Proven IAPA |  |  | V285-58 | Colonization |  |
| subtotal | 1 |  | 2 |  | 0 |  | 2 |  | 5 |
| **Resistant-no SNPs** |  |  |  |  |  |  |  |  |  |
| AmsterdamUMC |  |  | v215-17 |  | V265-17 |  | v284-29 |  | 3 |
| Radboudumc | v209-50 | Putative IA | v226-02 | ABPA, CCPA/CFPA | v273-30 | CPA | v293-59 | Colonization | 4 |
| subtotal | 1 |  | 2 |  | 2 |  | 2 |  |  |
| Total | 18 |  | 21 |  | 23 |  | 10 |  | 72 |

IPA; Invasive Pulmonary Aspergilloses, ABPA; allergic bronchopulmonary aspergillosis, IA; invasive aspergillosis, CNS; central nervous system, IAPA; Influenza‐associated pulmonary aspergillosis, CCPA; chronic cavity pulmonary aspergillosis, CFPA; chronic fibrosing pulmonary aspergillosis.

**Table S2.** *F_ST_* peaks

| *F_ST_* region | *F_ST_* Max | Chr | Genes | Putative gene function | Reference |
| --- | --- | --- | --- | --- | --- |
| 1 | 0.359534 | 1 | Afu1g03200 | mfsC Putative major facilitator superfamily (MFS) transporter |  |
|  |  |  | Afu1g03210 | Myb family transcription factor | González-Tobón J., Childers R. et al. “Searching for the Mechanism that Mediates Mefenoxam-Acquired Resistance in *Phytophthora infestans* and How It Is Regulated.” Phytopathology 2022 112(5): 1118-1133. Doi: 10.1094/PHYTO-07-21-0280-R. |
| 2 | 0.344901 | 1 | Afu1g11350 | MFS transporter of unknown specificity. Contains domain(s) with predicted role in transmembrane transport and integral component of membrane localization. | Del Sorbo, G., Schoonbeek, H.J. et al. “Fungal transporters involved in efflux of natural toxic compounds and fungicides.” Fungal genetics and biology (2000) 30(1): 1-15. Doi: 10.1006/fgbi.2000.1206. |
| 3 | 0.38147 | 1 | Afu1g14870 | gwt1 GPI-anchored wall transfer protein 1 | Hamoto M., Aizawa R. et al. “A novel fungicide aminopyrifen inhibits GWT-1 protein in glycosylphosphatidylinositol-anchor biosynthesis in Neurospora crassa.” Pesticide biochemistry and physiology 156 (2019): 1-8. Doi: 10.1016/j.pestbp.2019.02.013 |
| 4 | 0.444025 | 1 | Afu1g15050 | Hsp70 family chaperone | Blatzer M., Blum G. et al. “Blocking Hsp70 enhances the efficiency of amphotericin B treatment against resistant Aspergillus terreus strains.” Antimicrobial agents and chemotherapy 59.7 (2015): 3778-3788. Doi: 10.1128/AAC.05164-14.  Nagao J., Cho T. et al. “Candida albicans Msi3p, a homolog of the Saccharomyces cerevisiae Sse1p of the Hsp70 family, is involved in cell growth and fluconazole tolerance.” FEMS yeast research 12.6 (2012): 728-737. Doi: 10.1111/j.1567-1364.2012.00822.x |
| 5 | 0.332256 | 1 |  |  |  |
| 6 | 0.435706 | 1 | Afu1g16880 | ABC multidrug transporter | Del Sorbo, G., Schoonbeek, H.J. et al. “Fungal transporters involved in efflux of natural toxic compounds and fungicides.” Fungal genetics and biology (2000) 30(1): 1-15. Doi: 10.1006/fgbi.2000.1206. |
|  |  |  | Afu1g16910 | MFS multidrug transporter | Del Sorbo, G., Schoonbeek, H.J. et al. “Fungal transporters involved in efflux of natural toxic compounds and fungicides.” Fungal genetics and biology (2000) 30(1): 1-15. Doi: 10.1006/fgbi.2000.1206. |
| 7 | 0.329733 | 1 | Afu1g17530 | MFS transporter, putative | Del Sorbo, G., Schoonbeek, H.J. et al. “Fungal transporters involved in efflux of natural toxic compounds and fungicides.” Fungal genetics and biology (2000) 30(1): 1-15. Doi: 10.1006/fgbi.2000.1206. |
| 8 | 0.355542 | 2 |  |  |  |
| 9 | 0.426685 | 3 | Afu3g07810 | Succinate dehydrogenase flavoprotein subunit |  |
|  |  |  | Afu3g07870 | agd3 Galactosaminogalactan deacetylase |  |
|  |  |  | Afu3g07930 | Putative glutathione S-transferase | Cheng X., Dai T. et al. “Cytochrome P450 and Glutathione S-Transferase Confer Metabolic Resistance to SYP-14288 and Multi-Drug Resistance in Rhizoctonia solani.” Frontiers in microbiology 13 (2022): 806339-806339. Doi: 10.3389/fmicb.2022.806339 |
|  |  |  | Afu3g07950 | Contain domain(s) with predicted role in transmembrane transport and integral component of membrane localization |  |
| 10 | 0.356762 | 3 | Afu3g14670 | MFS transporter, putative | Del Sorbo, G., Schoonbeek, H.J. et al. “Fungal transporters involved in efflux of natural toxic compounds and fungicides.” Fungal genetics and biology (2000) 30(1): 1-15. Doi: 10.1006/fgbi.2000.1206. |
| 11 | 0.408914 | 4 | Afu4g03590 | MFS transporter, putative | Del Sorbo, G., Schoonbeek, H.J. et al. “Fungal transporters involved in efflux of natural toxic compounds and fungicides.” Fungal genetics and biology (2000) 30(1): 1-15. Doi: 10.1006/fgbi.2000.1206. |
| 12 | 0.448447 | 4 | Afu4g03740 | MFS transporter, putative | Del Sorbo, G., Schoonbeek, H.J. et al. “Fungal transporters involved in efflux of natural toxic compounds and fungicides.” Fungal genetics and biology (2000) 30(1): 1-15. Doi: 10.1006/fgbi.2000.1206. |
|  |  |  | Afu4g03750 | phthalate transporter, putative |  |
|  |  |  | Afu4g03800 | cytochrome P450 alkane hydroxylase, putative | Panwar S, Krishnamurthy S et al. “CaALK8, an alkane assimilating cytochrome P450, confers multidrug resistance when expressed in a hypersensitive strain of Candida albicans.” Yeast 18.12 (2001): 1117-1129. Doi: 10.1002/yea.762. |
| 13 | 0.378065 | 4 | Afu4g03920 | MFS drug transporter, putative | Del Sorbo, G., Schoonbeek, H.J. et al. “Fungal transporters involved in efflux of natural toxic compounds and fungicides.” Fungal genetics and biology (2000) 30(1): 1-15. Doi: 10.1006/fgbi.2000.1206. |
| 14 | 0.350669 | 4 | Afu4g04318 | copper resistance protein Crd2, putative |  |
| 15 | 0.396625 | 4 | Afu4g06180 | casein kinase II beta subunit CKB1 | Troppens D, Dmirtiev R, et al. “Genome-wide investigation of cellular targets and mode of action of the antifungal bacterial metabolite 2, 4-diacetylphloroglucinol in Saccharomyces cerevisiae.” FEMS yeast research 13.3 (2013): 322-334. Doi: 10.1111/1567-1364.12037. |
| 16 | 0.56444 | 4 | Afu4g07020 | mitochondrial cytochrome b2-like, putative |  |
| 17 | 0.53784 | 4 | Afu4g07210 | Mitochondrial acetolactate synthase small subunit |  |
|  |  |  | Afu4g07380 | Ortholog(s) have role in histone exchange and NuA4 histone acetyltransferase complex, Swr1 complex localization | William J., Loguinov A. et al. “Comparative functional genomic analysis identifies distinct and overlapping sets of genes required for resistance to monomethylarsonous acid (MMAIII) and arsenite (AsIII) in yeast.” Toxicological Sciences 111.2 (2009): 424-436. Doi: 10.1093/toxsci/kfp162. |
| 18 | 0.502988 | 4 |  |  |  |
| 19 | 0.380851 | 4 |  |  |  |
| 20 | 0.427191 | 4 | Afu4g08170 | succinate-semialdehyde dehydrogenase Uga2, putative | Ferreira A., Alves D., et al. “Synthesis of coumarin and homoisoflavonoid derivatives and analogs: The search for new antifungal agents.” Pharmaceuticals 15.6 (2022): 712. Doi: [10.3390/ph15060712](https://doi.org/10.3390%2Fph15060712) |
| 21 | 0.379459 | 4 |  |  |  |
| 22 | 0.447376 | 4 | Afu4g08740 | MFS multidrug transporter, putative | Del Sorbo, G., Schoonbeek, H.J. et al. “Fungal transporters involved in efflux of natural toxic compounds and fungicides.” Fungal genetics and biology (2000) 30(1): 1-15. Doi: 10.1006/fgbi.2000.1206. |
|  |  |  | Afu4g08800 | ABC a-pheromone efflux pump AtrD | Del Sorbo, G., Schoonbeek, H.J. et al. “Fungal transporters involved in efflux of natural toxic compounds and fungicides.” Fungal genetics and biology (2000) 30(1): 1-15. Doi: 10.1006/fgbi.2000.1206. |
|  |  |  | Afu4g09080 | C2H2 transcription factor (Seb1), putative, putative |  |
| 23 | 0.407882 | 5 |  |  |  |
| 24 | 0.404695 | 5 | Afu5g06060 | sulfur metabolism regulator SkpA, putative |  |
|  |  |  | Afu5g06070 | ABC multidrug transporter | Del Sorbo, G., Schoonbeek, H.J. et al. “Fungal transporters involved in efflux of natural toxic compounds and fungicides.” Fungal genetics and biology (2000) 30(1): 1-15. Doi: 10.1006/fgbi.2000.1206. |
| 25 | 0.441337 | 7 | Afu7g01490 | Major facilitator superfamily (MFS) peptide transporter | Del Sorbo, G., Schoonbeek, H.J. et al. “Fungal transporters involved in efflux of natural toxic compounds and fungicides.” Fungal genetics and biology (2000) 30(1): 1-15. Doi: 10.1006/fgbi.2000.1206. |
| 26 | 0.436339 | 7 | Afu7g01670 | MFS transporter Fmp42, putative | Del Sorbo, G., Schoonbeek, H.J. et al. “Fungal transporters involved in efflux of natural toxic compounds and fungicides.” Fungal genetics and biology (2000) 30(1): 1-15. Doi: 10.1006/fgbi.2000.1206. |
|  |  |  | Afu7g01740 | sugar transporter, putative |  |
|  |  |  | Afu7g01790 | MFS transporter, putative | Del Sorbo, G., Schoonbeek, H.J. et al. “Fungal transporters involved in efflux of natural toxic compounds and fungicides.” Fungal genetics and biology (2000) 30(1): 1-15. Doi: 10.1006/fgbi.2000.1206. |
| 27 | 0.408751 | 7 | Afu7g01970 | RTA1 domain protein, putative | Manente, M. Rta1, a yeast plasma membrane protein which confers resistance to fungicides. Diss. UCL-Université Catholique de Louvain, 2008. |
| 28 | 0.369973 | 7 | Afu7g05450 | SUN domain protein (Uth1), putative | Ritch J., Davidson S. et al. “The Saccharomyces SUN gene, UTH1, is involved in cell wall biogenesis.” FEMS yeast research 10.2 (2010): 168-176. Doi: 10.1111/j.1567-1364.2009.00601.x |
| 29 | 0.356575 | 7 | Afu7g05550 | sugar transporter family protein |  |
| 30 | 0.419127 | 7 | Afu7g05680 | siroheme synthase Met8 | Dietl A., Binder U. et al. “Siroheme is essential for assimilation of nitrate and sulfate as well as detoxification of nitric oxide but dispensable for murine virulence of Aspergillus fumigatus.” Frontiers in microbiology 9 (2018): 2615. Doi: 10.3389/fmicb.2018.02615 |
|  |  |  | Afu7g05700 | Ortholog(s) have ARF guanyl-nucleotide exchange factor activity and role in ER to Golgi vesicle-mediated transport, autophagosome assembly, cellular response to drug, hyphal growth, intra-Golgi vesicle-mediated transport |  |
|  |  |  | Afu7g05830 | Ortholog(s) have plasma membrane localization |  |
| 31 | 0.413253 | 8 | Afu8g00140 | MFS transporter, putative | Del Sorbo, G., Schoonbeek, H.J. et al. “Fungal transporters involved in efflux of natural toxic compounds and fungicides.” Fungal genetics and biology (2000) 30(1): 1-15. Doi: 10.1006/fgbi.2000.1206. |
| 32 | 0.355405 | 8 | Afu8g00770 | sugar transporter family protein |  |
| 33 | 0.399162 | 8 | Afu8g00885 | Ortholog(s) have plasma membrane localization |  |
|  |  |  | Afu8g00930 | Putative chitosanase |  |
|  |  |  | Afu8g00940 | MFS multidrug transporter, putative | Del Sorbo, G., Schoonbeek, H.J. et al. “Fungal transporters involved in efflux of natural toxic compounds and fungicides.” Fungal genetics and biology (2000) 30(1): 1-15. Doi: 10.1006/fgbi.2000.1206. |
| 34 | 0.42519 | 8 | Afu8g00962 | cytochrome P450, putative | Luo C., Schnabel G. “The cytochrome P450 lanosterol 14α-demethylase gene is a demethylation inhibitor fungicide resistance determinant in Monilinia fructicola field isolates from Georgia.” Applied and Environmental Microbiology 74.2 (2008): 359-366. Doi: 10.1128/AEM.02159-07 |
| 35 | 0.461115 | 8 | Afu8g01260 | Ergosteryl-3-O-glycine synthase ErgS |  |
|  |  |  | Afu8g01340 | Major facilitator superfamily (MFS) sugar transporter | Del Sorbo, G., Schoonbeek, H.J. et al. “Fungal transporters involved in efflux of natural toxic compounds and fungicides.” Fungal genetics and biology (2000) 30(1): 1-15. Doi: 10.1006/fgbi.2000.1206. |
| 36 | 0.409072 | 8 |  |  |  |

Sexual crossing was conducted to analyse the segregation patterns of azole phenotype and *cyp*51A haplotype. When one locus causes the azole resistant phenotype, according to Mendel’s law of segregation a 1:1 segregation in the offspring is expected when crossing an azole susceptible with a resistant isolate, and for two loci a 3:1 segregation. Six TR_34_/L98H and six TR_46_/Y121F/T289A *cyp*51A gene haplotypes azole resistant isolates were crossed to an azole sensitive *cyp*51A wild type isolate of opposing mating type. Among the offspring of ten out of twelve crosses the resistant to sensitive phenotype ratio was not significantly different from 1:1. This is indicative of triazole resistance being determined by a single locus. Crosses C028 and C029, which shared the same resistant parent (18A1), were the notable exception as they appeared to have 3:1 resistant to sensitive resistance ratio. This suggests that the resistant parent of this cross harboured a second independently segregating resistance locus. For the 10 crosses where resistance was most probably governed by a single locus, we also genotyped a random subset of 16 out of the 32 offspring. In all ten of these crosses the TR genotype was strongly linked with a triazole resistant phenotype. In the few cases were this was not the case are most likely the result of strong variations in mycelial growth rates present among the offspring. These variations in growth rates, in combination with our strict cut-off of a 1mm growth diameter on 8 mg/L itraconazole likely led to occasional misinterpretation when scoring when scoring the offspring for azole resistance.

**Table S3** Azole resistance segregation patterns of TR_34_/L98H and TR_46_/Y121F/T289A *cyp*51A gene haplotypes sexual crosses with a WT *cyp*51A gene haplotype.

|  | Azole phenotype | Cross number (Parent 1 x Parent 2) | | | | | | | | | | | | Sum |
| --- | --- | --- | --- | --- | --- | --- | --- | --- | --- | --- | --- | --- | --- | --- |
|  |  | C001 (88C19 X 46A23) | C003 (50C32 X 4A18) | C012  (16C32 X 38C31) | C013 (79C1 X 29C8) | C017  (46A23X 82C21) | C018 (46A23 X 88C19) | C019  (46A23 X  93C2) | C023 (5A16 X  93C2) | C027  (84C7 X  46A23) | C028  (18A1 X  Afir964) | C029  (18A1 X  93C2) | C031  (25C20 X  84C7) |  |
| Geno-phenotype counts  (N=16 per cross) | Res-TR | 9 | 10 | 6 | 6 | 6 | 8 | 8 | 5 | 8 | NA | NA | 8 | 74 |
|  | Res-WT | 0 | 1 | 0 | 0 | 0 | 0 | 0 | 1 | 0 | NA | NA | 2 | 4 |
|  | Sen-TR | 0 | 1 | 1 | 0 | 0 | 0 | 0 | 0 | 0 | NA | NA | 1 | 3 |
|  | Sen-WT | 7 | 4 | 9 | 10 | 10 | 8 | 8 | 10 | 8 | NA | NA | 5 | 79 |
| Phenotype counts  (N =32 per cross) | Resistant | 16 | 18 | 12 | 16 | 16 | 16 | 19 | 15 | 15 | 24 | 22 | 16 |  |
|  | Susceptible | 16 | 14 | 20 | 16 | 16 | 16 | 13 | 17 | 17 | 8 | 9 | 15 |  |
|  | Χ^2^ | 0 | 0.5 | 2 | 0 | 0 | 0 | 1.125 | 0.125 | 0.125 | 8 | 5.451613 | 0.032258 |  |
|  | p-value | 1 | 0.4795 | 0.157299 | 1 | 1 | 1 | 0.288844 | 0.723674 | 0.723674 | **0.004678** | **0.01955** | 0.857462 |  |

Χ^2^ Chi-square value

**Table S4** Microdilution antifungal susceptibility testing, in mg/L of 46 *A. fumigatus* isolates from this study. SNPs known to be correlated to fungicide resistance are highlighted in blue. MIC values of phenotypically fungicide resistance isolates are highlighted red, antifungal ECOFFs and clinical breakpoints for moulds were determined according the EUCAST E.Def 7.3, E.Def 9.4 and E.Def 11.0 procedures version 3.0 (ITZ/VOR 1 mg/L, POS 0·25 mg/L, ISA 2 mg/L), a wild type upper limit (WT-UL) was used for 0·25 mg/L for OLO (doi: 10.1128/AAC.00487-18) and 2 mg/L for TEB, DIF, PROT, PYR, AZOX, CARB, BEN (doi:10.1371/journal.pone.0031801). <https://www.eucast.org/astoffungi/clinicalbreakpointsforantifungals>

|  |  | target gene | medical azoles | | | | agricultural azoles | | | target gene | QoIs | | target gene | MBCs | | DHOH |
| --- | --- | --- | --- | --- | --- | --- | --- | --- | --- | --- | --- | --- | --- | --- | --- | --- |
| Isolate type | ID | **Cyp51A** | ITZ | VOR | POS | ISA | TEB | DIF | PROT | **CytB** | PYR | AZOX | **BenA** | CARB | BEN | OLO |
| Clinical | V259-78 | TR34 | >16 | 4 | 1 | 8 | 2 | 2 | <0·031 | G143A | 8 | >32 | F219Y | >16 | >32 | 0·063 |
| Clinical | V261-31 | TR34 | >16 | 16 | 2 | 8 | 2 | 8 | 0·031 | G143A | 4 | >32 | F219Y | >16 | >32 | 0·031 |
| Clinical | V204-54 | TR46 | >16 | >16 | 1 | >16 | >16 | 8 | 0·25 | G143A | >16 | >32 | F219Y | >16 | >32 | 0·063 |
| Clinical | V297-39 | TR46 | 0·5 | >16 | 0·25 | >8 | >16 | ND | ND | G143A | >16 | >32 | F219Y | >16 | >32 | 0·031 |
| Environmental | 11A6 | TR34 | >16 | 4 | 1 | 16 | 4 | 8 | 0·064 | G143A | 8 | 16 | F219Y | >16 | >32 | 0·063 |
| Environmental | 46A23 | TR34 | >16 | 16 | 1 | 8 | 2 | 4 | <0·031 | G143A | >16 | >32 | F219Y | >16 | >32 | 0·031 |
| Environmental | 60C3 | TR34 | ≥8 | >16 | 1 | >16 | >16 | >32 | >4 | G143A | 8 | 32 | F219Y | >16 | >32 | 0·063 |
| Environmental | 25C20 | TR46 | 1 | >16 | 0·5 | >16 | 16 | 2 | 0·031 | G143A | >16 | >32 | F219Y | >16 | >32 | 0·063 |
| Environmental | 66A16 | TR46 | 0·5 | >16 | 0·5 | >16 | 8 | 16 | 0·064 | G143A | >16 | >32 | F219Y | >16 | >32 | 0·031 |
| Clinical | V200-72 | TR34 | >16 | 4 | 1 | >16 | 1 | 2 | <0·031 | - | 0·25 | 1 | F219Y | >16 | >32 | 0·031 |
| Clinical | V199-25 | TR34 | >16 | 16 | 1 | 8 | 2 | 2 | 0·031 | - | 0·125 | 1 | F219Y | >16 | >32 | 0·031 |
| Clinical | V200-50 | TR34 | >16 | 16 | 1 | 8 | 4 | 4 | 0·031 | G134A | 2 | 16 | - | 1 | 1 | 0·031 |
| Clinical | V210-08 | TR34 | >16 | 8 | 1 | 8 | 4 | 8 | 0·031 | G134A | >16 | >32 | - | 1 | 2 | 0·063 |
| Clinical | V259-28 | TR34 | >16 | 4 | 1 | 4 | 2 | 2 | 0·064 | G134A | ≥16 | >32 | - | 1 | 2 | 0·031 |
| Environmental | 102C11 | TR34 | >16 | 2 | 1 | 16 | 2 | 2 | 0·064 | - | 0·125 | 1 | F219Y | 8 | 8 | 0·063 |
| Environmental | 40A21 | TR34 | >16 | 16 | 2 | 8 | 2 | 8 | 0·064 | - | 4 | >32 | F219Y | >16 | >32 | 0·031 |
| Environmental | 78C2 | TR34 | >16 | 4 | 0·5 | 8 | 4 | 4 | 0·031 | G134A | >16 | >32 | - | 1 | 2 | 0·016 |
| Environmental | 52B4 | TR46 | 0·5 | >16 | 0·5 | >16 | 8 | 8 | 0·064 | G134A | 8 | 32 | - | >16 | >32 | 0·031 |
| Environmental | 56C20 | TR46 | 0·5 | >16 | 0·5 | >16 | 16 | 16 | 0·25 | G134A | 4 | >32 | - | >16 | >32 | 0·063 |
| Clinical | V192-81 | TR34 | >16 | 4 | 0·5 | 8 | 2 | 1 | <0·031 | - | 0·125 | 0·5 | - | 1 | 2 | 0·031 |
| Clinical | V215-18 | TR34 | >16 | 8 | 2 | 8 | 1 | 8 | 0·064 | - | 0·125 | 0·25 | - | 1 | 2 | 0·063 |
| Clinical | V233-43 | TR34 | >16 | 4 | 0·5 | 8 | 0·5 | 4 | 0·064 | - | 0·016 | 0·125 | - | 0·5 | 2 | 0·063 |
| Clinical | V261-45 | TR34 | >16 | 8 | 1 | 8 | 4 | 4 | 0·064 | - | 0·063 | 0·5 | - | 1 | 1 | 0·063 |
| Clinical | V226-28 | TR46 | >16 | >16 | 1 | >16 | >16 | ND | ND | - | 0·125 | 1 | - | 1 | 2 | 0·063 |
| Environmental | 35CS28 | TR34 | >16 | 8 | 1 | 8 | 2 | 4 | <0·031 | - | 0·063 | 0·25 | - | 1 | 2 | 0·031 |
| Environmental | 4A19 | TR34 | >16 | 4 | 1 | 8 | 2 | 2 | <0·031 | - | 0·25 | 1 | - | 1 | 2 | 0·031 |
| Environmental | 9C23 | TR34 | >16 | 8 | 1 | 8 | 4 | 2 | 0·031 | - | 0·063 | 1 | - | >16 | >32 | 0·031 |
| Environmental | 48A5 | TR34 | >16 | 4 | 1 | 8 | 4 | 1 | 0·064 | - | 0·25 | 2 | - | 1 | 2 | 0·031 |
| Environmental | 82C23 | TR34 | >16 | 8 | 1 | 16 | 2 | 4 | 0·031 | - | 0·5 | 4 | - | 1 | 2 | 0·031 |
| Clinical | V226-02 | G54R | >16 | 2 | >2 | 2 | 0·5 | <0·25 | <0·031 | - | 0·031 | 0·063 | - | 1 | 2 | 0·031 |
| Clinical | V265-17 | wt | >16 | >16 | 1 | >16 | 0·5 | 8 | 0·125 | - | 0·063 | 0·063 | - | 1 | 2 | 0·063 |
| Clinical | V273-30 | wt | 16 | 0·5 | 8 | 1 | 1 | 0·25 | ≤0·031 | - | 0·063 | 0·25 | - | 1 | 2 | 0·016 |
| Clinical | V284-29 | wt | >16 | 4 | 1 | 4 | 4 | 8 | 0·031 | - | 0·25 | 1 | - | 0·125 | 2 | 0·031 |
| Environmental | 11A5 | wt | 0·25 | 0·5 | 0·064 | 0·5 | 0·5 | 0·25 | <0·031 | - | 0·25 | 1 | - | 1 | 2 | 0·031 |
| Environmental | 8C2 | wt | 1 | 2 | 0·25 | 2 | 2 | 1 | <0·031 | - | 0·063 | 0·063 | - | 1 | 2 | 0·031 |
| Environmental | 38C31 | wt | 0·25 | 1 | 0·064 | 1 | 0·5 | 0·25 | <0·031 | G134A | 0·063 | 4 | F219Y | 1 | 2 | 0·063 |
| Clinical | V197-79 | wt | 0·25 | 2 | 0·125 | 1 | 0·5 | 0·5 | <0·031 | - | 0·25 | 4 | - | 1 | 2 | 0·031 |
| Clinical | V208-52 | wt | 0·25 | 1 | 0·125 | 1 | 0·25 | 0·25 | 0·031 | - | 0·125 | 1 | - | 1 | 2 | 0·031 |
| Clinical | V231-63 | wt | 0·25 | 1 | 0·125 | 1 | 0·5 | 0·5 | <0·031 | - | 0·063 | 0·25 | - | 1 | 2 | 0·063 |
| Clinical | V234-13 | wt | 0·25 | 1 | 0·125 | 1 | 0·25 | 0·25 | <0·031 | - | 0·031 | 0·5 | - | 1 | 2 | 0·031 |
| Clinical | V254-55 | wt | 0·25 | 1 | 0·125 | 1 | 0·5 | ND | ND | - | 0·125 | 0·5 | - | 1 | 2 | 0·031 |
| Environmental | 11A2 | wt | 0·25 | 0·5 | 0·064 | 1 | 0·25 | 1 | <0·031 | - | 0·063 | 0·125 | - | 1 | 2 | 0·031 |
| Environmental | 55C1 | wt | 0·125 | 0·5 | 0·064 | 0·5 | 0·5 | 0·5 | <0·031 | - | 0·125 | 0·125 | - | 1 | 2 | 0·031 |
| Environmental | 78C1 | wt | 0·25 | 0·5 | 0·064 | 1 | 0·5 | 0·25 | <0·031 | - | 0·063 | 0·25 | - | 0·5 | 1 | 0·016 |
| Environmental | 82C21 | wt | 0·25 | 0·5 | 0·064 | 1 | 0·5 | 1 | <0·031 | - | 0·125 | 2 | - | 1 | 2 | 0·031 |
| Environmental | 98C11 | wt | 0·25 | 1 | 0·064 | 0·5 | 0·5 | 0·25 | <0·031 | - | 0·25 | 1 | - | 1 | 2 | 0·016 |

QoI; Quinone outside inhibitors, MBC; methyl benzimidazol-2-yl carbamate, ITZ; itraconazole, VOR; voriconazole, POS; posaconazole, ISA; isavuconazole, TEB; tebuconazole, DIF; difenoconazole, PROT; prothioconazole, PYR; pyraclostrobin, AZOX; azoxystrobin, CARB; carbendazim, BEN; benomyl, OLO; oloforim. ND: not determined.

**References:**

1.	Celia-Sanchez B, Mangum B, Gómez Londoño L, Wang C, Shuman B, Brewer M, et al. Pan-azole- and multi-fungicide-resistant *Aspergillus fumigatus* is widespread in the United States. Appl Environ Microbiol. 2024 Apr 17;90(4):e01782-23.

2.	Barber AE, Sae-Ong T, Kang K, Seelbinder B, Li J, Walther G, et al. Aspergillus fumigatus pan-genome analysis identifies genetic variants associated with human infection. Nat Microbiol. 2021 Nov 24;6(12):1526–36.

3.	Rhodes J, Abdolrasouli A, Dunne K, Sewell TR, Zhang Y, Ballard E, et al. Population genomics confirms acquisition of drug-resistant Aspergillus fumigatus infection by humans from the environment. Nat Microbiol. 2022 Apr 25;7(5):663–74.

4.	Kang SE, Sumabat LG, Melie T, Mangum B, Momany M, Brewer MT. Evidence for the agricultural origin of resistance to multiple antimicrobials in *Aspergillus fumigatus* , a fungal pathogen of humans. Heitman J, editor. G3 GenesGenomesGenetics. 2022 Feb 4;12(2):jkab427.

5.	Zhao S, Ge W, Watanabe A, Fortwendel JR, Gibbons JG. Genome-Wide Association for Itraconazole Sensitivity in Non-resistant Clinical Isolates of Aspergillus fumigatus. Front Fungal Biol. 2021 Jan 14;1:617338.

6.	Etienne KA, Berkow EL, Gade L, Nunnally N, Lockhart SR, Beer K, et al. Genomic Diversity of Azole-Resistant Aspergillus fumigatus in the United States. Cowen LE, editor. mBio. 2021 Aug 31;12(4):e01803-21.

7.	Lofgren LA, Lorch JM, Cramer RA, Blehert DS, Berlowski-Zier BM, Winzeler ME, et al. Avian-associated *Aspergillus fumigatus* displays broad phylogenetic distribution, no evidence for host specificity, and multiple genotypes within epizootic events. Sachs M, editor. G3 GenesGenomesGenetics. 2022 May 6;12(5):jkac075.

8.	Lofgren LA, Ross BS, Cramer RA, Stajich JE. The pan-genome of Aspergillus fumigatus provides a high-resolution view of its population structure revealing high levels of lineage-specific diversity driven by recombination. Sil A, editor. PLOS Biol. 2022 Nov 17;20(11):e3001890.

9.	Barber AE, Riedel J, Sae-Ong T, Kang K, Brabetz W, Panagiotou G, et al. Effects of Agricultural Fungicide Use on Aspergillus fumigatus Abundance, Antifungal Susceptibility, and Population Structure. Lorenz M, editor. mBio. 2020 Dec 22;11(6):e02213-20.

10.	Vasimuddin Md, Misra S, Li H, Aluru S. Efficient Architecture-Aware Acceleration of BWA-MEM for Multicore Systems. In: 2019 IEEE International Parallel and Distributed Processing Symposium (IPDPS). 2019. p. 314–24.

11.	Li H, Handsaker B, Wysoker A, Fennell T, Ruan J, Homer N, et al. The Sequence Alignment/Map format and SAMtools. Bioinformatics. 2009 Aug 15;25(16):2078–9.

12.	Auwera GAV der, O’Connor BD. Genomics in the Cloud: Using Docker, GATK, and WDL in Terra. O’Reilly Media, Incorporated; 2020. 496 p.

13.	Chang CC, Chow CC, Tellier LC, Vattikuti S, Purcell SM, Lee JJ. Second-generation PLINK: rising to the challenge of larger and richer datasets. GigaScience. 2015 Dec 1;4(1):s13742-015-0047–8.

14.	R Core Team. R: A Language and Environment for Statistical Computing [Internet]. Vienna, Austria: R Foundation for Statistical Computing; 2022. Available from: https://www.R-project.org/

15.	Wickham H, Averick M, Bryan J, Chang W, McGowan LD, François R, et al. Welcome to the Tidyverse. J Open Source Softw. 2019 Nov 21;4(43):1686.

16.	Huson DH, Bryant D. Application of Phylogenetic Networks in Evolutionary Studies. Mol Biol Evol. 2006 Feb 1;23(2):254–67.

17.	Ortiz EM. vcf2phylip v2.0: convert a VCF matrix into several matrix formats for phylogenetic analysis. [Internet]. [object Object]; 2019 [cited 2024 May 8]. Available from: https://zenodo.org/record/2540861

18.	Minh BQ, Schmidt HA, Chernomor O, Schrempf D, Woodhams MD, Von Haeseler A, et al. IQ-TREE 2: New Models and Efficient Methods for Phylogenetic Inference in the Genomic Era. Teeling E, editor. Mol Biol Evol. 2020 May 1;37(5):1530–4.

19.	Yu G, Smith DK, Zhu H, Guan Y, Lam TT. ggtree : an r package for visualization and annotation of phylogenetic trees with their covariates and other associated data. McInerny G, editor. Methods Ecol Evol. 2017 Jan;8(1):28–36.

20.	Danecek P, Auton A, Abecasis G, Albers CA, Banks E, DePristo MA, et al. The variant call format and VCFtools. Bioinformatics. 2011 Aug 1;27(15):2156–8.

21.	Buil JB, van der Lee H a. L, Rijs AJMM, Zoll J, Hovestadt J a. MF, Melchers WJG, et al. Single-Center Evaluation of an Agar-Based Screening for Azole Resistance in Aspergillus fumigatus by Using VIPcheck. Antimicrob Agents Chemother. 2017 Dec;61(12):e01250-17.

22.	Lex A, Gehlenborg N, Strobelt H, Vuillemot R, Pfister H. UpSet: Visualization of Intersecting Sets. IEEE Trans Vis Comput Graph. 2014 Dec 31;20(12):1983–92.

23.	Conway JR, Lex A, Gehlenborg N. UpSetR: an R package for the visualization of intersecting sets and their properties. Hancock J, editor. Bioinformatics. 2017 Sep 15;33(18):2938–40.

24.	Fraczek MG, Zhao C, Dineen L, Lebedinec R, Bowyer P, Bromley M, et al. Fast and Reliable PCR Amplification from *Aspergillus fumigatus* Spore Suspension Without Traditional DNA Extraction. Curr Protoc Microbiol [Internet]. 2019 Sep [cited 2023 Jul 28];54(1). Available from: https://onlinelibrary.wiley.com/doi/10.1002/cpmc.89

25.	Snelders E, van der Lee HAL, Kuijpers J, Rijs AJMM, Varga J, Samson RA, et al. Emergence of Azole Resistance in Aspergillus fumigatus and Spread of a Single Resistance Mechanism. Chris Kibbler, editor. PLoS Med. 2008 Nov 11;5(11):e219.

26.	Firke S. janitor: Simple Tools for Examining and	Cleaning Dirty Data. [Internet]. 2021. Available from: https://CRAN.R-project.org/package=janitor

27.	Wei, T S V. R package “corrplot”: Visualization of a Correlation Matrix. (Version 0.92) [Internet]. Available from: https://github.com/taiyun/corrplot.

28.	Wickham H. ggplot2: Elegant Graphics for Data Analysis. 2nd ed. 2016. Cham: Springer International Publishing : Imprint: Springer; 2016. 1 p. (Use R!).
